# Unlocking Programmable and Creative RNA Sequence Design with RDiffusion

**DOI:** 10.64898/2026.06.13.732023

**Authors:** Jue Wang, Jintong Dong, Lan Yang, Tianhao Li, Jianwei Yin, Jintao Chen, Stan Z. Li, Ying Dong, Jia Li, Cheng Tan

## Abstract

RNA, a pillar of the central dogma, has shaped three billion years of evolution. Despite cataloging tens of millions of non-coding RNAs and annotating millions, we have barely scratched the surface of the vast, unexplored RNA sequence space. Here, we introduce RDiffusion, a discrete-diffusion-based generative transformer model for RNA sequence design. Conditioned on diverse biological features, such as desired function, family type, secondary and tertiary structure, or binding protein partners, it can guide the generation of novel RNA sequences tailored to specific design specifications. We evaluate RDiffusion across a broad spectrum of RNA design tasks and find that it not only surpasses all baseline methods in design success rate and sequence diversity but also achieves advanced performance on downstream tasks. To demonstrate its translational potential, we applied RDiffusion with a customized seed-screening pipeline to de novo design therapeutic MicroRNA(miRNA) mimics in osteoarthritis (OA). This strategy identified mimic 0-4 as a lead candidate targeting SDC4 to preserve ECM homeostasis and cartilage integrity, with robust efficacy validated in clinical OA samples. Altogether, RDiffusion holds strong potential to serve as a cornerstone for RNA generation and representation learning, establishing a generalizable AI-driven paradigm for RNA therapeutic discovery.

## 1 Introduction

Over three billion years of evolution, DNA and proteins are recognized as the protagonists in molecular biology by most of people—DNA as the repository of genetic information and proteins as the executors of cellular functions. In the traditional “central dogma,” RNA is often relegated to the role of a mere messenger between these two [1, 2]. In reality, RNA transcends this intermediary function. It is the only biological macromolecule that can both store genetic information (e.g., mRNA and viral RNA) and act as a catalytic or regulatory molecule (e.g., ribozymes and non-coding RNAs), similar to proteins [3–5]. Consequently, RNA participates in nearly all aspects of life, and its engineering has become a powerful platform for developing therapeutics, diagnostics, and synthetic biological systems [6, 7].

In current RNA engineering, sequence design serves as the only direct entry point for altering RNA function, because the only entities that can be synthesized, modified, amplified, or sequenced in the laboratory are sequences composed of A, U, G, and C. Beyond serving as an effective tool for probing and understanding the physical principles of RNA folding, sequence design is also a core driving force in building synthetic biological systems and creating programmable therapeutics, diagnostics, and biosensors. It allows us to write “biological solutions” to specific problems in life sciences — much like programming a computer. Despite its potential, designing novel functional RNA sequences remains a formidable challenge [8, 9]. This complexity arises from RNA’s multi-level structural organization: its primary structure (linear nucleotide sequence), secondary structure (local helices, loops, and bulges formed by A-U and G-C base pairing), tertiary structure (complex three-dimensional shapes from long-range interactions), and even quaternary or quinary structures (supramolecular assemblies with other molecules or weak interactions with cellular metabolites) [10, 11]. Compounding this structural complexity is the vast functional diversity of RNA and the paucity of comprehensive annotations. Of all collected RNA sequences, only about 3% are cited in the literature, and merely 10,000 sequences have detailed functional annotations (e.g., Gene Ontology terms) [12–14]. The sequence space of possible RNAs is astronomically larger than the space of functional sequences. Furthermore, current annotations are heavily biased toward classical RNAs (e.g., rRNA and tRNA), leaving the functions of “dark matter” RNAs—such as long non-coding RNAs (lncRNAs) and circular RNAs (circRNAs)—largely unexplored [12–14]. Thus, the rational design of functional RNA remains a long and arduous endeavor.

The advent of artificial intelligence and the explosion in data scale offer unprecedented opportunities for RNA design [15, 16]. At the primary structure level, RNAcentral [17] provides the largest repository of non-coding RNA sequences, containing over 45 million entries. Leveraging this database, foundation models such as RNA-FM [18] and ERNIE-RNA [19] employ masked language modeling [20] to learn powerful sequence representations, significantly enhancing downstream RNA prediction tasks. For secondary structure, datasets like bpRNA-1m [21] and ArchiveII [22] have driven progress in both structure prediction and sequence design. Classical methods such as RNAFold [23] use dynamic programming based on nearest-neighbor thermodynamic models, while RFold [24] employs a Seq2map attention module with physical constraints to improve accuracy. RNAinformer [25] adopts a transformer [26] architecture that represents secondary structure directly as an adjacency matrix, enabling the modeling of complex features like non-canonical base pairs and pseudoknots. For tertiary structure, databases such as RNASolo [27, 28] provide high-quality, purified 3D structures. Models like RhoFold [29] integrate sequence features from RNA-FM with multiple sequence alignment (MSA) [30] information for accurate tertiary structure prediction. Meanwhile, RDesign [31] and gRNAde [32] leverage geometric graph neural networks [33] to model tertiary structure for design tasks. For functional design, GARDN [34] employs a generative adversarial network to generate sequences with desired functions, and RfamGen [35] uses a variational autoencoder [36] to sample novel functional RNA family [37] sequences from a continuous latent representation.

Despite these advances, existing methods are constrained by their reliance on single-condition inputs. RNA is a complex molecule with multi-level structures and diverse functions, and neglecting this multifaceted nature significantly limits performance and generalization [10, 11]. The grand challenge, therefore, is to harness all known constraints—from sequence and secondary structure to tertiary folds and functional annotations—to navigate the vast sequence space and create desired RNA molecules.

In this paper, we propose RDiffusion, a conditional diffusion-based model for comprehensive RNA sequence design. RDiffusion integrates diverse conditioning signals, including desirable functions, RNA family types, secondary structures, tertiary structures, binding protein information and so on, within a discrete diffusion framework [38, 39]. To support this model, we introduce RFuncData, a large-scale dataset comprising 11.5 million RNA sequences with associated functional and secondary structure annotations. After pretraining RDiffusion on RFuncData, we evaluate it across a series of design tasks: unconditional generation, functional design, secondary structure inverse design, tertiary structure inverse design, and binding-protein-based design(Section 2.2-2.6). RDiffusion consistently achieves state-of-the-art performance and high design success rates compared to specialized models in each domain. Meanwhile, we evaluate RDiffusion on multiple downstream prediction tasks alongside foundation models such as RNA-FM and ERNIE-RNA, and observe that RDiffusion surpasses these models as well, demonstrating that its pretraining paradigm offers a significant advance over existing representation learning methods. Furthermore, to validate the clinical value of RDiffusion, we deploy RDiffusion for the de novo design of functional miRNA sequences for Osteoarthritis disease(Section 2.7). The targeting efficacy and therapeutic potential of these designed candidates are subsequently validated through multiple biological experiments, establishing a pioneering paradigm for precision RNA therapeutics in joint diseases. We anticipate that RDiffusion will accelerate the programmable engineering of RNA—with profound implications for human health [40], drug development [41, 42], and gene-editing [43, 44] tools—while establishing a new standard for representation learning on RNA-related tasks.

## 2 Results

### 2.1 Overview of RDiffusion

Fig. 1 presents the overall workflow of RDiffusion, organized into six panels (a–f) that span data preparation, model architecture, generation process, and downstream applications. Fig. 1a illustrates the composition of the pretraining dataset, RFunc-Data, which comprises 11.5 million RNA sequences, each annotated with functional descriptions and secondary structures. Fig. 1b shows the diverse datasets used for fine-tuning across various downstream tasks. These datasets include RNA sequences paired with multiple condition types, including functional attributes (e.g., 5’UTR MRL [45, 46]), secondary structure, tertiary structure, RNA family classification, and binding proteins. Fig. 1c depicts the inference stage, where masked or partially masked RNA sequences, together with various conditioning signals, are input into the model. A guidance module is incorporated to steer the generation toward sequences with enhanced properties. Fig. 1d illustrates the forward and reverse processes of discrete diffusion. In the forward process, a raw RNA sequence is progressively masked token by token at each time step until all positions are masked. Conversely, in the reverse process, RDiffusion iteratively recovers each masked token to reconstruct the complete sequence. Fig. 1e presents the architecture of RDiffusion. The masked or partially masked RNA sequence is passed through an RNA embedding module to extract sequence features, which are subsequently refined by 12 transformer layers. Notably, the conditions at each stage are fed into the condition injector module as auxiliary information, enabling the RNA sequence design process to be constrained by user-defined specifications. RDiffusion iteratively recovers masked tokens under the reverse process until the sequence is fully reconstructed. Furthermore, during the inference stage, a guidance model evaluates the quality of the generated sequence at each time step and provides guidance via gradient descent to improve the generation process. Finally, Fig. 1f lists several design tasks to which RDiffusion can be applied, including unconditional, functional, structure-based, and binding-based RNA design, among others.

**Fig. 1.**
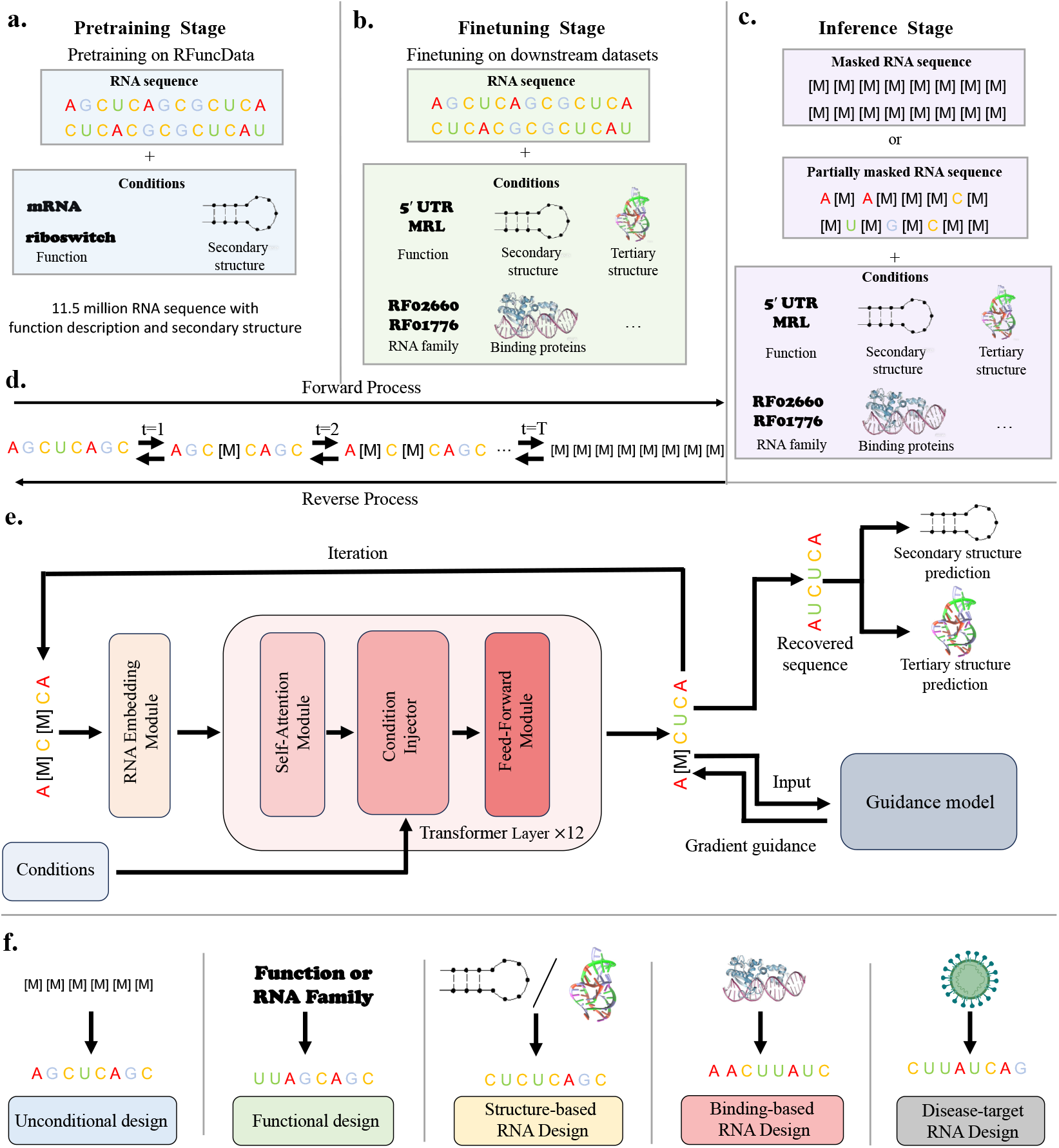
The overall workflow of RDiffusion. **a** Pretraining data composition; **b** Fine-tuning data composition across diverse downstream tasks; **c** Inference-stage inputs and the guidance model for conditional generation; **d** Forward and reverse processes of discrete diffusion; **e** Model architecture of RDiffusion; **f** Representative downstream applications enabled by RDiffusion.

### 2.2 Functional RNA Design with RDiffusion

The exploitation of RNA functionality has enabled the development of diverse synthetic molecular systems, with profound implications for basic research, biomanufacturing [47], and medical applications [48]. However, experimental screening for RNA sequences with desired functions remains costly and inefficient, primarily owing to the vast sequence space that expands exponentially with sequence length. Consequently, the development of computational methods capable of accurate and diverse RNA sequence design according to specified functional requirements is both essential and challenging. To address this, we apply RDiffusion to three representative functional RNA design tasks: RNA family classification, 5’UTR mean ribosome loading (MRL) optimization, and the Target RNA activity of guide RNA (gRNA) [49, 50], as elaborated in the following sections.

#### 2.2.1 Function: RNA Family Type

To investigate RDiffusion’s performance in functional RNA design, we first apply it to the Rfam [37] dataset to generate novel RNA sequences conditioned on a specific RNA family type. Rfam provides a coherent classification framework that organizes functionally diverse RNA molecules into families based on evolutionary origins and functional characteristics, thereby revealing the hierarchical and networked nature of biological regulation. The dataset comprises 320,477 sequences spanning 3,395 RNA family types, each annotated with secondary structure and family label. We randomly split the data into training (80%), validation (10%), and test (10%) sets.

Fig. 2a illustrates the workflow of RDiffusion for functional RNA design. A fully masked RNA sequence is input into RDiffusion, with the functional condition serving as guidance. At each time step, the partially generated sequence is fed into a guidance model—implemented as either a regression or classification model—to predict the target functional property. The resulting prediction loss (mean squared error for regression or cross-entropy for classification) is computed, and gradient backpropagation from the guidance model provides additional gradient information to steer the generation toward functionally desirable sequences.

**Fig. 2.**
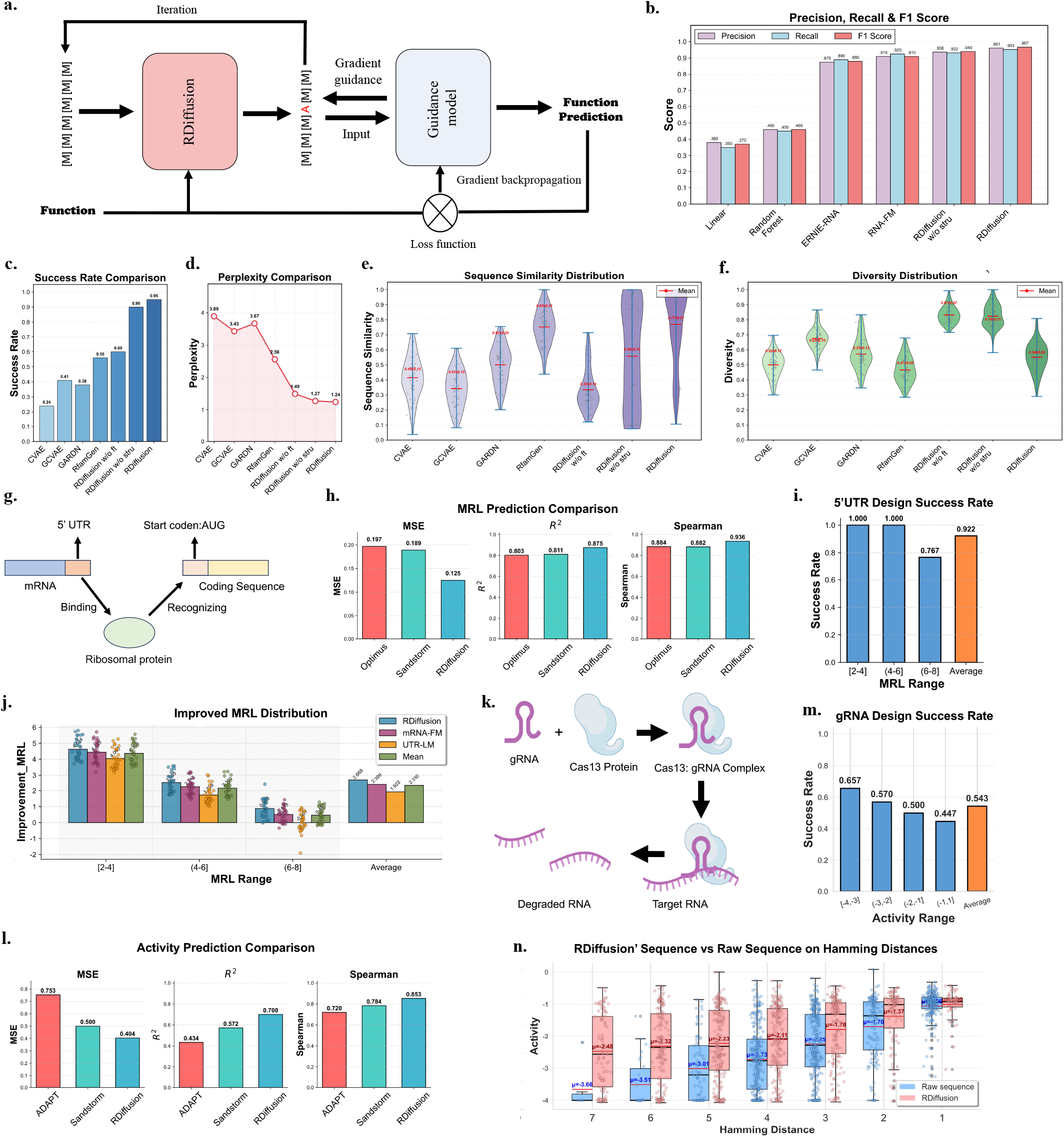
Functional RNA Design of RDiffusion based on RNA family, 5’UTR MRL and gRNA-Cas13 activity tasks. **a** The workflow of RDiffusion on RNA family design task. **b** The classification performance of 6 methods on the Rfam dataset with 3 metrics; **c** The success rate comparison of 7 methods; **d** The perplexity comparison of 7 methods; **e** The sequence similarity distribution of 7 methods; **f** The diversity distribution of 7 methods. “w/o ft” and “w/o stru” represent without finetuning and without structure. **g** Schematic illustration of mRNA 5’UTR functionality; **h** Comparison of three methods for 5’UTR MRL prediction (lower MSE indicates better performance); **i** Design success rate of 5’UTR across multiple MRL ranges; **d** Distribution of improved MRL across multiple MRL ranges; **j** Schematic illustration of gRNA and Cas13 protein functionality; **k** Comparison of three methods for predicting Cas13 binding activity to target RNA; **m** Design success rate of gRNA across multiple activity ranges; **n** Activity comparison between RDiffusion-generated sequences and raw sequences across seven Hamming distance bins.

Fig. 2b evaluates the generalization capability of RDiffusion via fine-tuning on an RNA family classification task, using Precision, Recall, and F1 score as evaluation metrics. We compare RDiffusion against four baselines—Linear classifier, Random Forest [51], ERNIE-RNA, and RNA-FM—under identical settings. For fair comparison, we also fine-tune a variant of RDiffusion without secondary structure input, as competing methods cannot incorporate such information. Classical machine learning methods (Linear and Random Forest) achieve overall scores below 0.5, indicating the non-trivial nature of this classification task. Among all methods, RDiffusion achieves the best performance across all three metrics: Precision 0.961, Recall 0.953, and F1 score 0.967. Even without secondary structure information, RDiffusion attains a Precision of 0.936, Recall of 0.932, and F1 score of 0.940, surpassing RNA-FM (Precision 0.910, Recall 0.925, F1 0.910) and ERNIE-RNA (Precision 0.875, Recall 0.890, F1 0.880). These results demonstrate the superior generalization capability and advanced pretraining strategy of RDiffusion.

Fig. 2c evaluates RDiffusion on RNA sequence generation conditioned on RNA family type, using design success rate as the metric. Success is defined as a sequence being correctly classified as belonging to the target RNA family. For a comprehensive evaluation, we employ three independent classifiers—RDiffusion, RNA-FM, and ERNIE-RNA—as evaluation tools, and a sequence is considered successful only if all three classifiers agree on its family assignment. We employ the RDiffusion classifier from Fig. 2b—which achieves the highest classification accuracy—as the guidance model. We evaluate RDiffusion under three settings: fine-tuned, fine-tuned without secondary structure input, and without fine-tuning. We compare against four baselines: CVAE [52], GCVAE [53], GARDN [34], and RfamGen [35], all trained on the same 8:1:1 training/validation/test split. In the evaluation stage, given a target RNA family label, models generate RNA sequences optionally conditioned on secondary structure. The fine-tuned RDiffusion achieves a leading success rate of 0.95, substantially outperforming all baselines. Notably, even without secondary structure input and relying solely on the family type prompt, RDiffusion attains a success rate of 0.90, far exceeding RfamGen (0.60), GARDN (0.38), GCVAE (0.41), and CVAE (0.24) (RfamGen and GARDN can accept secondary structure input). Remarkably, even the non-fine-tuned RDiffusion—which cannot accept secondary structure or recognize family labels and relies solely on guidance model steering—achieves a success rate of 0.60, surpassing all baselines. These results confirm that RDiffusion accurately generates novel RNA sequences conditioned on target RNA family types.

Fig. 2d presents perplexity scores of generated sequences across models. Perplexity reflects model confidence; lower perplexity indicates greater confidence. RDiffusion under all three settings achieves significantly lower perplexity than baselines, indicating that RDiffusion better captures the biological patterns embedded in RNA sequences and thus produces more confident predictions.

Fig. 2e and f depict sequence similarity and diversity distributions. Sequence similarity measures the degree of similarity between generated sequences and target family representatives, while diversity [54] quantifies the variability among generated sequences. RDiffusion achieves high average sequence similarity (0.77) but relatively low diversity (0.55), slightly outperforming RfamGen. This behavior reflects RDiffusion’s sensitivity to structural constraints imposed by secondary structure, which can be mitigated by adjusting the generation temperature. In contrast, the non-fine-tuned RDiffusion achieves the lowest sequence similarity (0.33) and the highest diversity (0.87) while maintaining a reasonably high success rate (0.60). Notably, the fine-tuned RDiffusion without structural constraints achieves a favorable balance among success rate (0.90), sequence similarity (0.56), and diversity (0.76). Collectively, these results demonstrate that RDiffusion not only accurately generates RNA sequences for desired family types but also produces novel and diverse sequences.

#### 2.2.2 Function: 5’UTR Sequence Mean Ribosomal Loading

Designing 5’UTR sequences with high MRL is of great significance for optimizing protein expression, as higher MRL generally indicates more efficient translation initiation and elongation, which is critical for biopharmaceutical production and synthetic biology applications [49, 50]. However, achieving this design goal presents substantial challenges. The primary difficulty lies in the complex and still poorly understood regulatory logic of the 5’UTR, where translation efficiency is governed by multiple overlapping signals—including secondary structures, upstream open reading frames (uORFs), and specific sequence motifs such as the Kozak context—all of which must be balanced to avoid unintended consequences such as cryptic promoters or RNA instability [55, 56]. Fig. 2g illustrates how the 5’UTR of an mRNA exerts its functionality. Through interactions with ribosomal proteins, the 5’UTR ensures proper positioning of the ribosome at the start codon, thereby governing translation initiation efficiency, a process essential for post-transcriptional gene expression control.

Following Optimus [57] and Sandstorm [34], we first applied RDiffusion to the 5’UTR MRL prediction task using the benchmark dataset from Optimus 5-prime [57], which comprises 83,919 artificially synthesized random 5’UTRs and 7,600 native human 5’UTRs, each with corresponding MRL values. As shown in Fig. 2h, RDiffusion substantially outperforms existing methods across all metrics: RDiffusion achieves an MSE of 0.125, an R^2^ of 0.875, and a Spearman correlation of 0.936; Sandstorm achieves 0.189, 0.811, and 0.882; Optimus achieves 0.197, 0.803, and 0.884, respectively. These results demonstrate RDiffusion’s strong generalization ability for detecting functional features in RNA sequences.

After fine-tuning RDiffusion for MRL prediction, we leveraged the pretrained RDiffusion as the policy model and the RDiffusion MRL predictor as the reward model to perform Group Reward Policy Optimization (GRPO) [58], yielding a new version of RDiffusion capable of designing novel 5’UTR sequences with high MRL. During inference, we randomly mask a UTR sequence (with mask ratios ranging from 0.1 to 0.9) and iteratively recover the entire sequence. Success is defined as a designed 5’UTR sequence having a higher predicted MRL than the raw sequence. To ensure accurate evaluation, we employed RDiffusion, mRNA-FM [18], and UTR-LM [59] to predict the MRL of designed sequences, and used the average of the three predictions as the final result. To comprehensively evaluate RDiffusion’s design performance, we randomly selected 30 5’UTR sequences from each of three MRL ranges: 2–4, 4–6, and 6–8. Fig. 2i presents the design success rates of RDiffusion. RDiffusion achieves a high average success rate of 0.922 across all MRL ranges, indicating that it has effectively learned the regulatory logic of 5’UTRs and can identify specific functional motifs. For sequences in the 2–4 and 4–6 ranges, the success rates reach 1.0. Notably, the success rate for the 6–8 range is relatively lower, which can be attributed to the fact that an MRL of 6–8 represents a very high baseline, making optimization sub-stantially more difficult than for lower-MRL sequences. Nevertheless, RDiffusion still attains a success rate of 0.767, demonstrating its robust design effectiveness. Fig. 2j depicts the distribution of MRL improvements across multiple MRL ranges. For raw 5’UTR sequences with low MRL, the improvements achieved by RDiffusion are substantial: the average improved MRL values are 4.366 for the 2–4 range and 2.160 for the 4–6 range, with all three predictor models reporting improvements above zero. For the high-MRL (6–8) range, the mean improvement remains positive, reaching 0.463. Overall, the average improved MRL across all sequences is 2.669, 2.399, and 1.922 for the three predictor models, with a mean of 2.330, collectively demonstrating clear and consistent improvement.

#### 2.2.3 Function: Target RNA Activity of the gRNA-Cas13 Ribonucleoprotein Complex

Unlike the well-known DNA-targeting Cas9 [50] system, CRISPR-Cas13 [60, 61] represents a class of RNA-guided CRISPR systems that specifically recognize and cleave single-stranded RNA. The guide RNA (gRNA) serves as the core determinant of Cas13 targeting specificity, functioning essentially as a programmable navigation system. Upon association with crRNA (gRNA), the Cas13 protein forms a ribonucleoprotein complex that scans cellular RNAs for sequences complementary to the gRNA. Upon encountering a fully matched target RNA, the complex becomes activated, leading to specific cleavage of the bound target RNA as well as indiscriminate degradation of surrounding single-stranded RNAs(Fig. 2k). In contrast to the previous task, where the primary design objective was simply to achieve high binding activity to the target RNA—typically attainable with fully complementary gRNAs—the current goal is to design gRNAs that simultaneously exhibit both high target-binding activity and enhanced target specificity. This dual requirement is critical to ensure precise targeting by Cas13 while avoiding partial complementarity to off-target transcripts, thereby minimizing off-target effects. Given that Cas13, once activated, elicits non-specific collateral cleavage, any off-target recognition could lead to unnecessary cytotoxicity. Consequently, the major challenge lies in navigating the vast sequence space to identify gRNAs that strike an optimal balance between high on-target activity and robust specificity [60, 61].

Analogous to the MRL prediction task, we fine-tuned RDiffusion on the ADAPT benchmark dataset [62], which comprises approximately 19,000 guide RNA–target pairs, to perform gRNA activity prediction. As shown in Fig. 2l, RDiffusion substantially outperforms existing methods across all metrics: RDiffusion achieves an MSE of 0.404, an R^2^ of 0.700, and a Spearman correlation of 0.853; Sandstorm achieves 0.500, 0.572, and 0.784; Optimus achieves 0.753, 0.434, and 0.720, respectively. These results further demonstrate the generalization capability of RDiffusion in detecting functional features of RNA sequences.

Following the same GRPO training and evaluation framework used for the 5’UTR design task, we further assessed the performance of RDiffusion on designing gRNAs with high activity and specificity. Success is defined as the generated sequence having a higher predicted activity than the raw sequence. Additionally, we randomly selected 300 guide–target RNA pairs across four activity ranges: [-4,-3], (-3,-2], (-2,-1], and (-1,0].

Fig. 2m presents the activity-based design results across these ranges. The average success rate across all ranges is 0.543, with the highest success rate of 0.657 observed in the [-4,-3] range. Success rates gradually decline as the baseline activity range increases. Nevertheless, even the lowest success rate, observed in the (-1,0] range, remains at 0.447, indicating effective and reliable design performance.

Fig. 2n plots the activity comparison between RDiffusion-generated sequences and raw sequences across different Hamming distances^1^. Across all Hamming distances, the average activity of generated sequences exceeds that of raw sequences, with particularly pronounced improvements at larger distances (Hamming distance 4–7). For instance, the average activity of generated sequences at Hamming distance 7 (-2.48) surpasses that of raw sequences at Hamming distance 4 (-2.73). More notably, as Hamming distance increases from 1 to 7, the raw sequences exhibit a clear monotonic decline in activity, dropping from -1.06 at distance 1 to -3.66 at distance 7, indicating that sequences farther from the optimal target are intrinsically less active. In contrast, RDiffusion-generated sequences maintain consistently higher activity across all distance levels (ranging from -1.00 to -2.48), demonstrating RDiffusion’s strong capability to recover high activity from sequences that are highly diverged from the optimal template. These results collectively demonstrate that RDiffusion is not only capable of designing sequences with high activity and high specificity but also capable of achieving higher activity within defined specificity constraints.

### 2.3 Secondary Structure-based RNA Design with RDiffusion

RNA secondary structure refers to the spatial conformation formed by intramolecular base pairing (primarily A-U, G-C, and weaker G-U pairs), resulting in local helical stems and unpaired loops [63]. It serves as the critical link between linear primary sequences and complex three-dimensional folding and constitutes the direct functional platform for most non-coding RNAs [63, 64]. Designing RNA sequences based on secondary structure fundamentally transforms the principle of “structure dictates function” from passive interpretation to active creation. Since RNA functions—such as molecular recognition, catalysis, and regulation—depend on specific three-dimensional motifs (e.g., stem-loops and pseudoknots) rather than on linear sequence alone, directly using a target secondary structure as a blueprint enables the encoding of sequences that autonomously fold and execute predetermined functions. This structure-guided design strategy holds substantial application potential [65, 66].

To evaluate RDiffusion’s design capability on structure-based tasks, we applied it to the bpRNA-1m dataset [21] and tested it on the bpRNA TS0 dataset. bpRNA-1m is one of the largest and most comprehensive meta-databases of RNA secondary structures, comprising 102,318 RNA sequences with their corresponding secondary structures. It integrates data from seven major databases, including Rfam, PDB [67], and CRW [68]. The TS0 test set is a standardized subset curated from bpRNA-1m by removing sequences with more than 80% similarity, resulting in 1,305 entries. Widely adopted as a benchmark test set by models such as SPOT-RNA [69] and ProtRNA [70], TS0 is designed to mitigate data leakage and ensure fair evaluation.

Fig. 3a illustrates the workflow of applying RDiffusion to secondary structure-based design. The procedure is analogous to the function-based design task: a masked sequence, together with the target secondary structure, is input into RDiffusion, which iteratively predicts masked tokens. A guidance model provides additional gradient information to refine the outputs. Furthermore, the generated sequences are fed into RibonanzaNet [71], an evaluator that computes OpenKnot and Eterna scores to assess computational prediction quality and potential synthesizability.

**Fig. 3.**
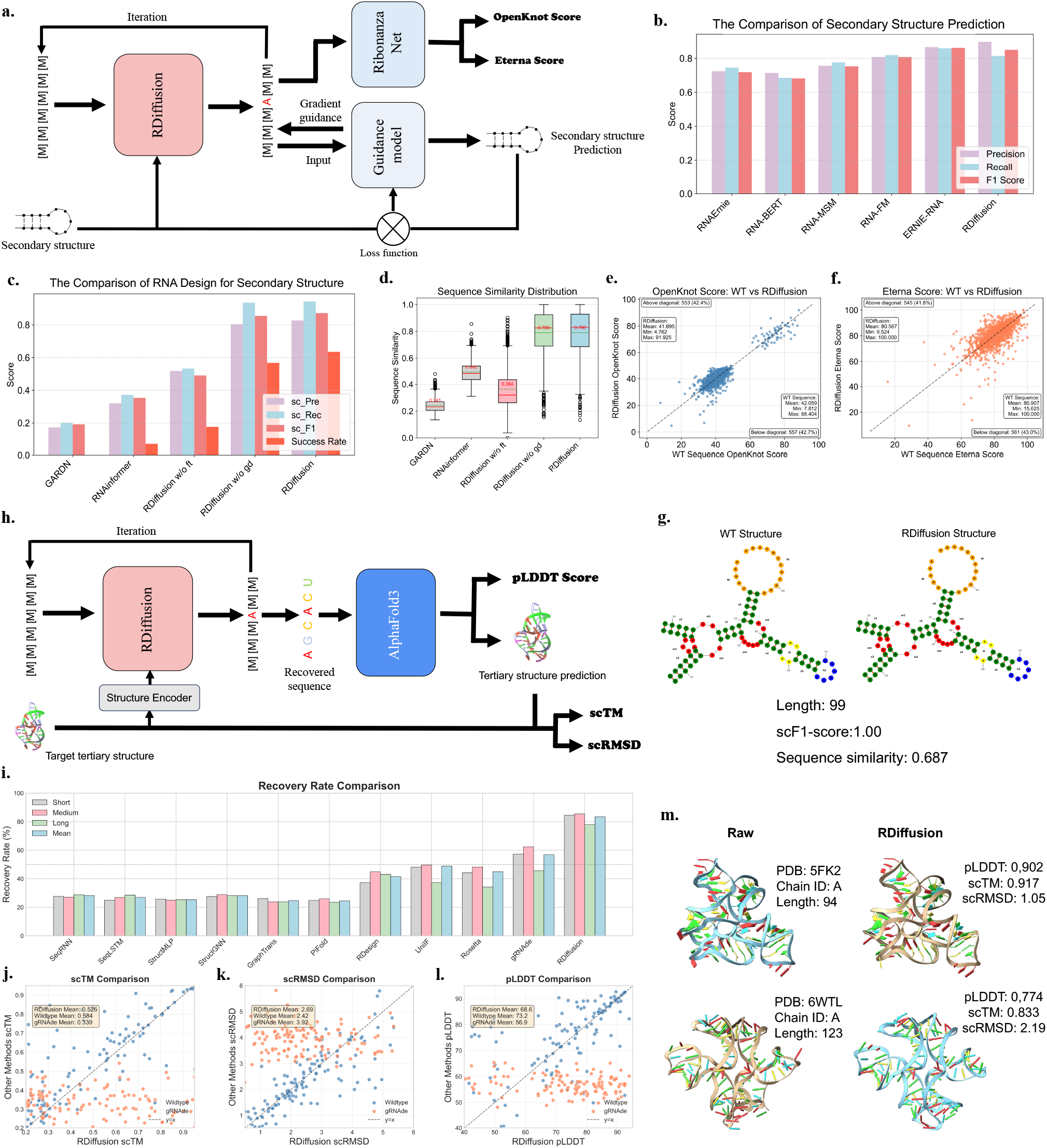
Secondary and Tertiary structure based RNA design of RDiffusion. **a** Workflow of RDiffusion for secondary-structure-based RNA design; **b** Secondary structure prediction performance of six methods on the TS80 dataset evaluated by three metrics; **c** RNA design task comparison of five models: scPre, scF1, and scRec denote self-consistency Precision, F1 score, and Recall, respectively; w/o gd indicates without guidance; **d** Comparison of sequence similarity distributions across five methods; **e** OpenKnot score comparison between RDiffusion-generated and wild-type sequences; **f** Eterna score comparison; **g** Visualization of exemplary secondary structures for wild-type and RDiffusion-designed sequences. **h** Workflow of RDiffusion for tertiary-structure-based RNA design; **i** Comparison of recovery rates across eleven methods; **j** Comparison of self-consistency TM (scTM) scores between RDiffusion and other methods; **k** Comparison of self-consistency RMSD (scRMSD) scores between RDiffusion and other methods; **l** Comparison of pLDDT scores between RDiffusion and other methods; **m** Two visualization examples of AlphaFold3-predicted structures based on sequences generated by RDiffusion.

Fig. 3b presents the secondary structure prediction results. We fine-tuned RDiffusion on the bpRNA-1m TR0 dataset for secondary structure prediction and evaluated its predictive performance on the TS0 dataset using three metrics. For fair comparison, we fine-tuned several baselines—ERNIE-RNA, RNA-FM, RNA-MSM, RNA-BERT, and RNAErnie—under identical settings on the TR0 dataset. RDiffusion achieves strong results (Precision 0.897, Recall 0.814, F1 score 0.850), comparable to ERNIE-RNA (Precision 0.866, Recall 0.859, F1 score 0.862) and superior to other baselines. Notably, RDiffusion attains the highest Precision among all models without using structural information leveraged by ERNIE-RNA, demonstrating its advanced generalization capability.

Fig. 3c evaluates RDiffusion on secondary structure-based RNA design using the bpRNA-1m dataset, with design performance assessed on the TS0 dataset via four metrics: self-consistency Precision (scPre), self-consistency Recall (scRec), self-consistency F1 score (scF1), and success rate. Self-consistency metrics are defined as the Precision, Recall, and F1 score between the target structure and the structure obtained by refolding the designed sequence. Success is defined as a folding model correctly folding a generated sequence into the target secondary structure. We employed ERNIE-RNA—trained on bpRNA-1m and RNAStralign [72] datasets—as both the folding model and the guidance model, as it achieves state-of-the-art secondary structure prediction performance. We evaluated RDiffusion under three settings: without fine-tuning on bpRNA-1m, without guidance model, and with fine-tuning on bpRNA-1m. Baselines include GARDN and RNAinformer, both trained on bpRNA-1m. As shown in Fig. 3c, both baselines achieve poor results (GARDN: scPre 0.173, scRec 0.201, scF1 0.192, success rate 0.0; RNAinformer: 0.320, 0.372, 0.354, 0.07), reflecting the high heterogeneity between TS0 and the bpRNA-1m training set and thus the difficulty of the task. In contrast, RDiffusion achieves high performance across all metrics (0.827, 0.943, 0.872, and 0.635). Notably, even without fine-tuning on bpRNA-1m, RDiffusion substantially outperforms all baselines (0.518, 0.533, 0.491, 0.176), demonstrating its robust design capability on unseen datasets. The version without guidance yields slightly lower results (0.804, 0.936, 0.855, 0.568), indicating that while the guidance model contributes positively, it is not essential for RDiffusion’s strong performance.

Fig. 3d shows the sequence similarity distributions of five models. Both RDiffusion and its version without guidance achieve relatively high average sequence similarity (0.79), reflecting their ability to capture sequence–structure relationships. Surprisingly, the fine-tuned RDiffusion exhibits a substantially lower average sequence similarity (0.364) than RNAinformer, yet still achieves high self-consistency and success rates, indicating that RDiffusion can discover novel and plausible sequences beyond the training distribution.

Fig. 3e and f compare OpenKnot and Eterna scores between wild-type sequences and RDiffusion-designed sequences. The OpenKnot Score and Eterna Score are two core metrics for evaluating RNA sequence design quality, both originating from the Eterna platform. The OpenKnot Score computationally predicts a sequence’s secondary structure using RNA folding algorithms and measures the base-pair distance or matching degree between the predicted and target structures; a higher score indicates greater consistency. The Eterna Score is typically obtained through experimental validation (here predicted by RibonanzaNet) using SHAPE chemical probing; a higher score indicates that the sequence folds more closely to the target structure under real-world conditions. RDiffusion-generated sequences achieve average OpenKnot and Eterna scores of 41.895 and 80.567, respectively, comparable to wild-type sequences (42.059 and 80.907). Notably, the maximum OpenKnot score of RDiffusion-designed sequences reaches 91.925, exceeding the best wild-type score of 88.404, demonstrating that RDiffusion can generate sequences that better conform to target structures than some naturally occurring sequences.

Fig. 3g provides a visualization example of folding results for a wild-type sequence and an RDiffusion-designed sequence. Using the ERNIE-RNA pretrained model to fold the RDiffusion sequence, we observe that the RDiffusion sequence completely adopts the wild-type structure despite having only 0.687 sequence similarity. This example demonstrates that RDiffusion generates structurally faithful RNA sequences that are genuinely novel, rather than merely memorizing and reproducing training set sequences.

### 2.4 Tertiary Structure-based RNA Design with RDiffusion

Designing RNA sequences based on their tertiary structures holds significant application value in fields such as synthetic biology, RNA drug development, and nanotechnology [73]. For example, engineering RNA molecules that fold into specific three-dimensional conformations enables gene expression regulation, biosensor construction, and targeted delivery vehicle development, while also facilitating the understanding of functional mechanisms in natural RNA systems. However, this task faces substantial difficulties: the free energy landscape of RNA folding is extremely complex, the mapping between sequence and tertiary structure remains incompletely understood, and existing prediction algorithms suffer from limitations in both accuracy and computational efficiency [31, 32]. Moreover, designed sequences may deviate from the target structure in cellular environments due to misfolding, non-specific molecular interactions, or environmental factors, while experimental screening and validation remain time-consuming and labor-intensive.

To address this significant challenge, we applied RDiffusion to the RNASolo dataset [27, 28], a comprehensive RNA tertiary structure database curated from the PDB containing 12,914 structures. Following the same preprocessing protocol as gRNAde, we retained structures with resolution 4.0 Å, resulting in 4,223 unique RNA structures from RNASolo. This dataset was split into 3,823 structures for training and 200 each for validation and testing.

Fig. 3h illustrates the tertiary-structure-based RNA design workflow of RDiffusion. The target tertiary structure is first passed through a structure encoder that extracts geometric information and compresses it into compact feature representations, which are then fed into RDiffusion together with masked RNA sequences. After sequence generation, we employ AlphaFold3 [29] to predict the corresponding tertiary structures of the generated sequences and obtain pLDDT [74–76] scores to measure prediction confidence, which also serves as an indicator of sequence validity. Furthermore, we compute the TM-score [77] and RMSD [78] between the target tertiary structure and the AlphaFold3 prediction to obtain self-consistency TM (scTM) and self-consistency RMSD (scRMSD) scores, assessing whether the designed sequences can refold into the target structures.

Fig. 3i presents the recovery rate comparison across eleven methods. Recovery rate measures the accuracy between raw RNA sequences and model-designed sequences, reflecting whether models can precisely inverse-design target tertiary structures. We compared RDiffusion with ten baselines, evaluating performance across RNA sequences of different lengths (short: 1–20 nt, medium: 21–200 nt, long: >200 nt) for comprehensive assessment. RDiffusion achieves substantially superior recovery rates, with an average of 0.83 and length-specific rates all exceeding 0.79. In contrast, all baselines achieve average recovery rates below 0.57, with the best results for short, medium, and long sequences being 57.34%, 62.35%, and 45.59%, respectively—substantially lower than RDiffusion. These results demonstrate that RDiffusion effectively captures the complex geometric information embedded in tertiary structures and successfully maps geometric features to sequence features.

Fig. 3j–l compare scTM, scRMSD, and pLDDT scores among RDiffusion-generated sequences, gRNAde-generated sequences, and wild-type sequences. RDiffusion consistently outperforms gRNAde across all three metrics: RDiffusion achieves average scTM of 0.526, scRMSD of 2.69 Å, and pLDDT of 68.6, compared to gRNAde’s 0.339, 3.92 Å, and 56.9, respectively. These results indicate that RDiffusion not only designs sequences more similar to the raw sequences but also produces sequences with superior structural rationality and alignment with the target structure. Furthermore, compared to wild-type sequences (scTM: 0.584, scRMSD: 2.42 Å, pLDDT: 73.2), RDiffusion-designed sequences achieve only slightly lower average results— particularly for scRMSD (2.42 Å vs. 2.62 Å)—and some designed sequences even outperform their wild-type counterparts.

Fig. 3m visualizes two examples of AlphaFold3-predicted structures for RDiffusion-designed sequences alongside the corresponding raw structures. For PDB 5FK2 (Chain A, length 94), the RDiffusion-designed sequence achieves a predicted pLDDT of 0.902, an scTM of 0.917, and a low scRMSD of 1.05 Å. For PDB 6WTL (Chain A, length 123), the RDiffusion-designed sequence yields a pLDDT of 0.774, an scTM of 0.833, and an scRMSD of 2.19 Å. These results demonstrate excellent structural rationality and alignment with the target tertiary structures.

### 2.5 Binding protein-based RNA Design with RDiffusion

Designing RNA sequences capable of specifically binding to a given protein represents a central endeavor in RNA biology, chemical biology, and synthetic biology [79, 80]. At its core, this approach endows RNA molecules with antibody-like recognition functions, enabling the programmable integration of protein-binding capabilities into RNA. This facilitates dynamic and reversible intervention in cellular functions, thereby providing a powerful molecular tool for deciphering regulatory mechanisms of life and for developing next-generation precision therapeutics [79, 80].

However, RNA design for specific protein binding is non-trivial. It not only involves the highly complex underlying grammar of RNA-protein interactions but also necessitates consideration of the inherent trade-off between high affinity and high specificity: an RNA that binds tightly to its intended protein target may also exhibit off-target binding to other unintended proteins due to structural similarity, thereby substantially increasing the difficulty of rational design.

To assess the versatility of RDiffusion’s design capability, we applied it to binding-protein-based RNA design tasks. Fig. 4a illustrates the workflow of RDiffusion for binding protein-based RNA design. A fully masked sequence is input into RDiffusion together with the specific binding protein names and icSHAPE values [81]. At each time step, a guidance model evaluates the binding affinity of the partially generated sequence to the target protein and provides gradient-based guidance to steer generation toward more desirable sequences.

**Fig. 4.**
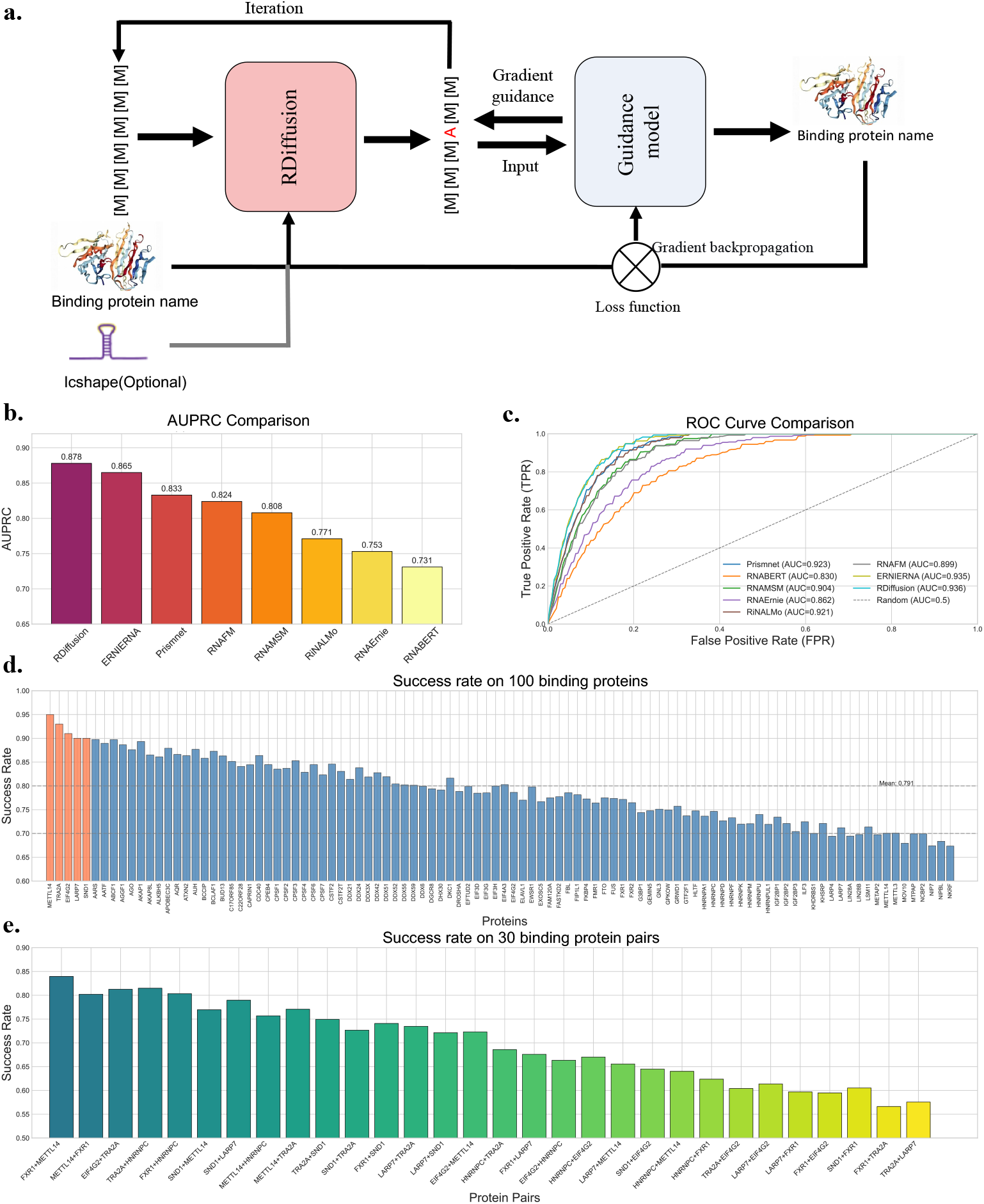
Binding Protein-based RNA Design of RDiffusion. **a** Workflow of RDiffusion for binding protein-based RNA design; **b** Comparison of AUPRC scores across eight methods; **c** Comparison of ROC curves across eight methods; **d** Design success rate of RDiffusion across 100 binding proteins; **e** Design success rate of RDiffusion across 30 binding protein pairs.

We conducted experiments using the benchmark dataset from PrismNet [82], which includes icSHAPE data. The dataset was divided into several subsets according to different RNA-binding proteins (RBPs) and cellular environments. We first employed RDiffusion to perform RNA-protein binding prediction, where the model must predict whether an RNA binds to a specific protein. Following the same setting as ERNIE-RNA, we selected 17 RBPs in the HeLa cell environment and compared against baselines including ERNIE-RNA, PrismNet, RNA-FM, RNA-MSM [83], RiNALMo [84], RNAErnie [85], and RNABert [86]. The fine-tuned RDiffusion on this prediction task was subsequently used as the guidance model.

Fig. 4b presents the AUPRC comparison of eight methods. RDiffusion achieves the best performance among all methods, with an AUPRC of 0.878. Furthermore, we plot the ROC curves of these methods(Fig. 4c) and observe that RDiffusion maintains stable accuracy across all true positive rates, demonstrating that RDiffusion effectively captures the hidden relationships governing RNA-protein interactions.

In Fig. 4d, we present the RNA design success rate of RDiffusion across 100 distinct binding proteins. For each target binding protein, we generate 100 candidate RNA sequences. A sequence is considered successfully designed if three independent models—RDiffusion, ERNIE-RNA, and PrismNet—all predict that it binds to the intended target protein. RDiffusion achieves consistently high success rates across all 100 proteins, with all rates exceeding 0.65. Notably, the top five binding proteins achieve success rates above 0.90.

Furthermore, to evaluate whether RDiffusion can design RNA sequences capable of binding to multiple proteins simultaneously, we selected 30 protein pairs and employed RDiffusion to generate sequences targeting each pair. For each protein pair, we generated 100 sequences and again used the three models for validation. As Fig. 4e shown, RDiffusion maintains robust performance under this more challenging setting, with success rates across all pairs exceeding 0.57 and the highest reaching 0.84. These results demonstrate that RDiffusion possesses not only strong capability for designing RNA sequences that bind to single target proteins but also the ability to generate sequences with specificity for multiple protein targets.

### 2.6 Unconditional RNA Design with RDiffusion

To validate whether RDiffusion can design reasonable and well-behaved RNA sequences, we applied it to an unconditional design task, generating RNA sequences of 20, 50, 100, 200, and 500 nucleotides, with 50 samples per length. In Extended Data Fig. 1a, to assess the quality of these generated sequences, we used RibonanzaNet to predict their secondary structures and calculated OpenKnot and Eterna scores between each sequence and its predicted secondary structure to evaluate structural stability. Furthermore, we employed AlphaFold3 to predict the tertiary structures of the generated sequences and obtained pLDDT scores, which reflect AlphaFold3’s per-residue prediction confidence. Generally, regions with pLDDT > 70 are predicted to have correct backbone conformations, while regions with pLDDT > 90 approach experimental structure accuracy. If a newly designed RNA sequence achieves consistently high pLDDT scores across its entire length (e.g., average > 70 or > 80) in AlphaFold3 predictions, it indicates that the sequence can fold into a stable, reasonable three-dimensional structure without obvious unfavorable features (such as long disordered regions, clashes, or unreasonable bends). Conversely, low pLDDT regions suggest that parts of the sequence may be overly flexible, poorly designed, or in need of further optimization. Thus, pLDDT serves as a quantitative metric for assessing the quality of generated RNA sequences.

Extended Data Fig. 1b and c present the OpenKnot and Eterna score distributions of RNA sequences generated by RDiffusion across five lengths. The average OpenKnot and Eterna scores for short sequences (20, 50, and 100 nt) are 40.8 and 81.6, respectively, maintaining high and stable performance comparable to wild-type sequences reported in Section 2.3 (OpenKnot: 42.1, Eterna: 80.9). The OpenKnot and Eterna scores range from 20.4 to 50.0 and from 40.8 to 100.0, respectively, further indicating that RDiffusion’s unconditional generation capability is stable and achieves a high success rate. In contrast, the long sequences generated by RDiffusion exhibit relatively lower performance, with average OpenKnot and Eterna scores of 25.2 and 50.3, respectively. This phenomenon arises primarily because long RNA sequences possess more complex structures and larger search spaces, which current secondary structure prediction models struggle to model effectively, leading to increased false-positive pairings.

Extended Data Fig. 1d shows the diversity distribution across generated sequences for the five lengths. RDiffusion achieves a substantial average diversity of at least 0.753 for length 500, with sequences of length 20 reaching 0.948. Furthermore, we observe that average diversity decreases as sequence length increases. This is because longer RNA sequences tend to contain more repetitive segments, which are often functionally required; in contrast, many short RNA sequences may deliberately avoid sequence repeats due to functional constraints. These results further demonstrate that RDiffusion not only achieves high design accuracy but also maintains a strong capacity for generating novel sequences.

Extended Data Fig. 1e illustrates two AlphaFold3-predicted tertiary structures for RNA sequences designed by RDiffusion. Both structures exhibit high pLDDT scores, ranging from 47.9 to 92.1 and from 53.6 to 83.4, with average pLDDT values of 74.10 and 74.21, respectively. These results further demonstrate that the sequences generated by RDiffusion exhibit good structural rationality, resulting in high pLDDT scores.

### 2.7 RDiffusion Enables De Novo Design of Therapeutic miRNAs for Osteoarthritis

miRNAs are a class of endogenous, small non-coding single-stranded RNA molecules that play a pivotal role in post-transcriptional gene regulation and exhibit significant correlations with the pathogenesis of various diseases [87, 88]. Osteoarthritis(OA) is a highly prevalent degenerative joint disease in which miRNAs play pivotal roles in orchestrating the pathological progression [89, 90]. Despite its severe clinical burden, current management remains largely confined to symptomatic relief, leaving an urgent unmet medical need for disease-modifying osteoarthritis drugs (DMOADs) [91]. Remarkably, the feasibility of localized intra-articular injection renders OA an ideal and highly translatable model for nucleic acid therapeutics [92].

To address these challenges, we employed RDiffusion to de novo design functional miRNA sequences targeting OA. First, we compiled a miRNA dataset from RNADisease [93] and retrieved the corresponding sequences from miRBase [94], with each entry including a sequence and its associated disease relevance score. We then fine-tuned RDiffusion on these miRNA sequences to perform a discrete diffusion task, enabling the model to learn the general functional patterns of miRNAs. Additionally, we fine-tuned RDiffusion to carry out weighted classification based on miRNA sequences and their disease relevance scores, thereby capturing disease-related patterns and establishing a guidance model. The output probability from this guidance model for OA is subsequently used to assess the relevance of a given miRNA sequence to the disease.

We thus established a robust pipeline that combines database mining, RDiffusion-based generative design, and systematic experimental evaluations to screen and validate promising OA-associated candidates. First, experimentally validated OA-associated miRNA sequences and their corresponding disease relevance scores were extracted from the RNADisease database and cross-referenced with miRBase. These miRNAs were subsequently clustered into distinct groups based on their seed region (nucleotides 2–8), allowing a maximum threshold of two mismatched nucleotides. To quantify the disease relevance of each unique seed, a weighted seed score was calculated using a custom scoring strategy where weighting factors were determined by the degree of sequence identity: perfectly identical sequences were assigned a full weight of 1, whereas sequences harboring one or two nucleotide mismatches were down-weighted to 6/7 and 5/7, respectively, during weighted average calculations. Seeds were strictly prioritized and selected for downstream computational design if they met any of the following multi-parametric criteria: a seed score greater than 0.95, a contributor count of at least 10 sequences, or a seed score greater than 0.90 accompanied by at least 5 contributor sequences. This rigorous screening workflow ultimately yielded 46 core candidate seeds. Using these 46 selected core seeds as input templates, the generative diffusion model RDiffusion was deployed to perform de novo design of novel OA-related miRNA sequences by introducing 0 to 3 nucleotide mutations within the seed regions. The newly generated variants were functionally and structurally evaluated using both the RDiffusion disease score, which reflects the probability of OA association, and the eterna score, which assesses RNA secondary structural stability. Within each seed-mutated group, sequences were ranked concurrently by these two metrics, and the top 20 candidates were selected. To delineate their downstream mechanisms, these optimized sequences were subjected to target prediction via the miRDB database, and the predicted targets were subsequently analyzed for functional enrichment using Metascape. Candidates demonstrating strong functional enrichment in critical OA-pathological pathways, such as cellular senescence, extracellular matrix (ECM) degradation, and chronic inflammation, were selected for chemical synthesis as double-stranded mimics. Finally, these synthetic mimics were transfected into chondrocytes and patient-derived cartilage tissues to systematically evaluate their regulatory phenotypes, functional outcomes, and targeting specificity in the context of joint degeneration. The detailed workflow is illustrated in Fig. 5a.

**Fig. 5.**
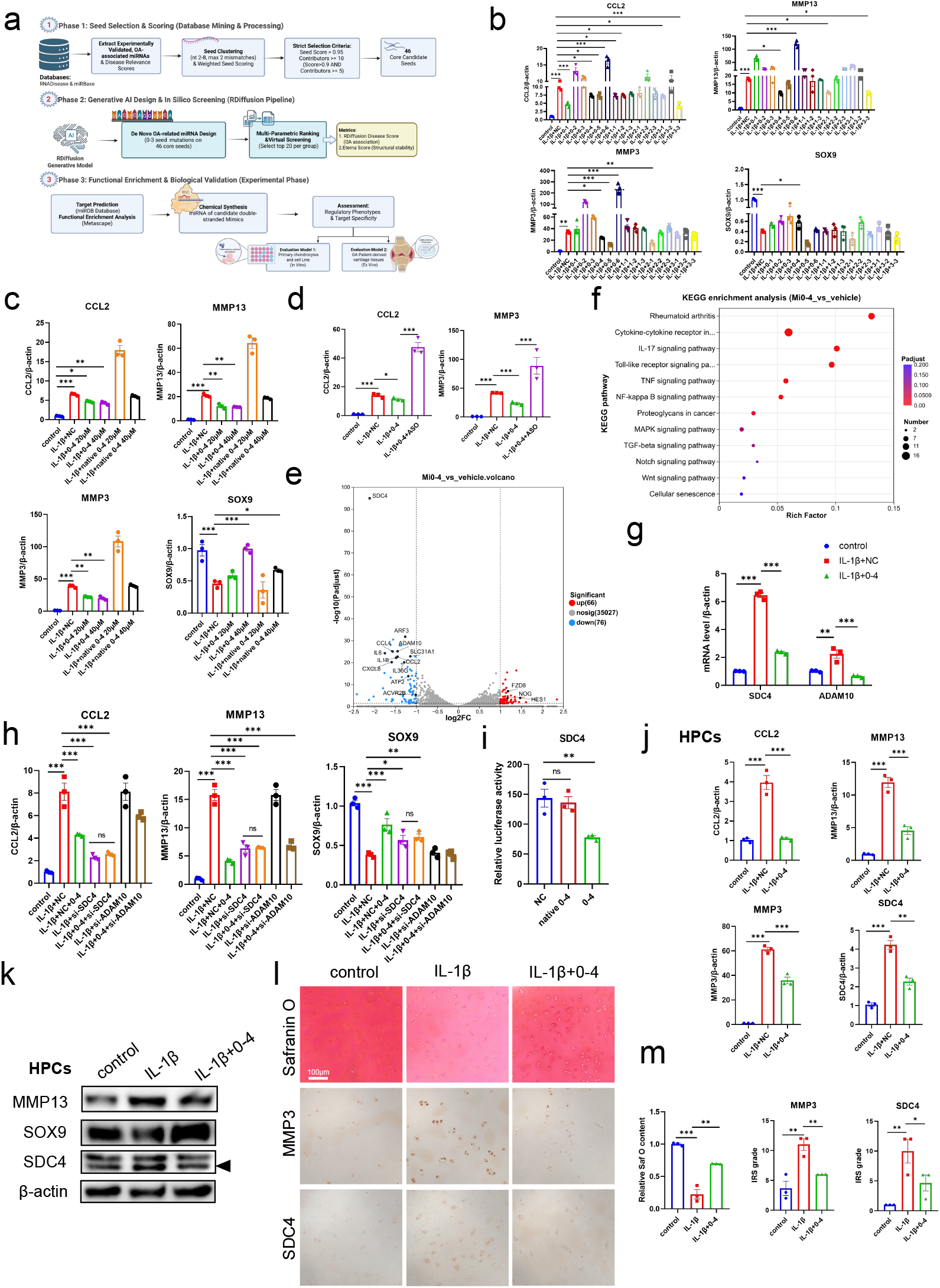
De novo RDiffusion-designed miRNA 0-4 attenuates osteoarthritis by suppressing SDC4. **a** The workflow of Osteoarthritis-target miRNA de novo design with RDiffusion and biological validation. **b** SW1353 cells were transfected with an equal amount of mimic (40 nM) for 24 h, followed by IL-1*β* stimulation for another 24 h. RT–qPCR analysis of ECM-related anabolic and catabolic genes and inflammatory cytokines. **c** RT–qPCR analysis of ECM-related genes and inflammatory cytokines in SW1353 cells transfected with increasing doses of mimic 0-4 or the native miRNA prior to IL-1*β* stimulation. d. RT–qPCR analysis of CCL2 and MMP3 expression in SW1353 cells transfected with mimic 0-4 alone or in combination with its ASO before IL-1*β* stimulation. e. Volcano plot showing differentially expressed genes between the vehicle + IL-1*β* group and the mimic 0-4 + IL-1*β* (Mi0-4) group. **f** KEGG pathway enrichment analysis of differentially expressed genes identified between the vehicle + IL-1*β* and Mi0-4 groups. **g** RT–qPCR validation of SDC4 and ADAM10 in SW1353 cells transfected with mimic 0–4 prior to IL-1*β* stimulation. **h** SW1353 cells were transfected with mimic 0-4, si-SDC4, or si-ADAM10 alone, or co-transfected with mimic 0-4 plus si-SDC4 or si-ADAM10, followed by IL-1*β* stimulation. ECM-related gene expression was assessed by RT–qPCR. **i** SDC4 3’-UTR dual-luciferase reporter assay in HEK293T cells co-transfected with reporter plasmids and mimic 0-4 or the corresponding native miRNA. Firefly luciferase activity was normalized to Renilla luciferase activity. **j** RT–qPCR analysis of ECM anabolic and catabolic markers in human primary chondrocytes (HPCs) transfected with mimic 0-4 and subsequently stimulated with IL-1*β*. **k** Immunoblot analysis of ECM-related proteins and SDC4 expression in HPCs treated as in (h). l-m. Representative images and quantitative analysis of Safranin O staining and immunohistochemistry staining for MMP3 and SDC4 in human cartilage explants. Data are shown as mean *±* SEM. ^*\**^*p <* 0.05, ^*\*\**^*p <* 0.01, ^*\*\*\**^*p <* 0.01.

Based on the selection criteria described above, 15 candidate miRNA mimics were synthesized and subjected to in vitro efficacy evaluation. In OA, dysregulation of the extracellular matrix (ECM), characterized by matrix metalloproteinase (MMP)-mediated degradation overcoming SOX9-driven ECM synthesis, serves as a primary driver of cartilage destruction, accompanied by persistent low-grade chronic inflammation [95, 96]. We first assessed these 15 mimics in an in-vitro chondrocyte model established by IL-1*β*-induced ECM dysregulation in SW1353 cells. As shown in Fig. 5b, seven mimics (e.g., 0-1, 0-4, and 0-5) markedly reduced CCL2 mRNA levels, displaying broad anti-inflammatory potential. Among these seven candidates, three (0-4, 2-1, and 3-3) also substantially reduced the expression of ECM catabolic factors MMP3 and MMP13. Considering the alterations in the ECM synthesis-related transcription factor SOX9, mimic 0-4 was uniquely capable of not only suppressing inflammation and ECM catabolism, but also promoting ECM anabolism. Conversely, mimic 0-6 showed an inverse pattern, driving the expression of CCL2, MMP3 and MMP13 while repressing SOX9. These preliminary findings suggest that mimic 0-4 exerts superior chondroprotective efficacy by preserving chondrocyte homeostasis, whereas mimic 0-6 disrupts it.

Since the seed regions of mimic 0-4 and 0-6 were not mutated, we hypothesized that the design of the non-seed sequences by RDiffusion might optimize target binding affinity and structural stability. To validate this hypothesis, we selected representative native miRNA (hsa-miR-140-3p for 0-4 and hsa-miR-9-3p for 0-6) sharing the same seed region for phenotypic comparison. As shown in Fig. 5c and Extended Data Fig. 2a, 0-4 exhibited significantly superior efficacy of ECM dysregulation compared to hsa-miR-140-3p. In contrast, mimic 0-6 displayed effects comparable to hsa-miR-9-3p, suggesting that the regulation of exacerbating ECM breakdown remains predominantly seed-driven. Consequently, we focused our subsequent investigations on the therapeutic efficacy of mimic 0-4.

During OA progression, senescent chondrocytes drive ECM dysregulation by secreting senescence-associated secretory phenotype (SASP) factors that accelerate collagen and aggrecan cleavage [97, 98]. Furthermore, we examined the anti-senescent potential of mimic 0-4 using an ionizing radiation (IR)-induced senescence model in SW1353 cells. Treatment with mimic 0-4 prominently protected against IR-induced senescence, markedly reducing the proportion of senescence-associated *β*-galactosidase (SA-*β*-gal)-positive cells and downregulated the cell cycle arrest marker p21 as well as the catabolic enzyme MMP3 (Extended Data Fig. 2b-c).

To confirm the target and sequence specificity of mimic 0-4, we synthesized its complementary antisense oligonucleotide (ASO). Notably, co-transfection of mimic 0-4 with its corresponding ASO substantially reversed the 0-4-mediated chondroprotective efficacy (Fig. 5d), demonstrating that the biological activity of mimic 0-4 is sequence-and target-specific. Further to systematically elucidate the molecular mechanisms and potential target by which the mimic 0-4 ameliorates ECM degradation and chondrocyte degeneration, we performed RNA-seq on IL-1*β*-stimulated chondrocytes treated with either mimic 0-4 or vehicle control. As shown in the hierarchical clustering analysis (Extended Data Fig. 2d), the biological replicates of the mimic 0-4-treated group and the vehicle group clustered into two highly distinct and homogeneous branches. Differential expression analysis identified a robust transcriptional remodeling, yielding a total of 142 differentially expressed genes (DEGs) that comprised 66 significantly up-regulated and 76 significantly down-regulated transcripts (Fig. 5e). Specifically, the down-regulation of key candidate chondrocyte regulators, such as SDC4, ADAM10, ACVR2B, and ATF2, was accompanied by the robust suppression of inflammatory cytokines (IL1*β*, IL6, CXCL8, and IL36G) and critical ECM catabolic effectors (notably MMP13). Conversely, the upregulation of protective factors such as NOG, HES1, and FZD8 activated anabolic and survival networks, which are crucial for preventing chondrocyte hypertrophy and cellular senescence. Consistently, KEGG pathway enrichment analysis revealed that the DEGs were pre-dominantly mapped to classical pathways driving osteoarthritis pathogenesis (Fig. 5f). Key enriched cascades included critical inflammatory networks (Rheumatoid arthritis, Cytokine-cytokine receptor interaction, IL-17, TNF, and NF-*κ*B signaling) and core regulators of chondrocyte fate (MAPK, Notch, Wnt, TGF-beta signaling, and Cellular senescence), demonstrating the potent chondroprotective and anti-inflammatory regulatory network of mimic 0-4.

Further analysis of the direct targets of mimic 0-4 was performed by cross-referencing the 76 significantly downregulated DEGs with the top 200 predicted targets of native miRNAs sharing the identical seed sequence retrieved from the miRDB database. Venn diagram analysis revealed a highly conserved intersection of 12 candidate genes (Extended Data Fig. 2e-f). We subsequently validated the expression levels of the 12 overlapping genes in chondrocytes subjected to IL-1*β* stimulation followed by mimic 0-4 treatment. Notably, among the upstream regulatory candidates, SDC4 (Syndecan-4) and ADAM10 (Disintegrin and metalloproteinase domain-containing protein 10) exhibited expression patterns that were highly consistent with our RNA-seq data (Fig. 5g, Extended Data Fig. 2g). Under IL-1*β* conditions, both SDC4 and ADAM10 were notably upregulated, while their aberrant elevations were robustly suppressed upon mimic 0-4 intervention. To determine whether mimic 0-4 mediated attenuation of ECM dysregulation depends on SDC4 or ADAM10, we performed siRNA-mediated knockdown of either gene, alone or in combination with mimic 0-4 transfection. As depicted in Fig. 5h and Extended Data Fig. S5h, the protective effect of mimic 0-4 on ECM homeostasis was largely abolished by SDC4 deficiency, but remained unaffected by ADAM10 deficiency, as indicated by changes in MMP13, CCL2, and SOX9 expression. SDC4 is a transmembrane heparan sulfate proteoglycan that regulates ECM interactions and cartilage homeostasis [99]. In OA cartilage, SDC4 is markedly upregulated and promotes extracellular matrix degradation by directly activating the catabolic aggrecanase ADAMTS5, consistently, SDC4 deficiency substantially mitigates cartilage destruction in murine OA models, making SDC4 a promising chondroprotective target [100, 101]. Next, we conducted dualluciferase reporter assays to confirm whether mimic 0-4 directly and specifically targets SDC4. Transfection of mimic 0-4 into HEK293T cells expressing a SDC4 3’-UTR luciferase reporter markedly reduced the luciferase activity, whereas the corresponding native miRNA (hsa-miR-140-3p) mimic exerted no discernible effect (Fig. 5i), confirming the specific targeting of the SDC4 3’-UTR by mimic 0-4. Consequently, non-seed sequence optimization by RDiffusion significantly enhanced target binding affinity toward SDC4, leading to its effective downregulation and the subsequent preservation of ECM homeostasis and cartilage integrity.

To enhance the clinical relevance of our findings, human primary chondrocytes (HPCs) and cartilage explants were isolated from OA patients undergoing total knee arthroplasty. In IL-1*β*-stimulated HPCs, mimic 0-4 also exerted a significant protective effect on ECM homeostasis, as evidenced by the reduced expression of MMP13, MMP3, and CCL2 (Fig. 5j). Notably, mimic 0-4 markedly suppressed SDC4 expression at both the transcriptional and protein levels (Fig. 5j-k). Furthermore, in the ex-vivo cartilage explants, mimic 0-4 significantly attenuated proteoglycan loss, as evidenced by Safranin O staining (Fig. 5l-m). Consistently, immunohistochemical (IHC) analysis showed that mimic 0-4 markedly reduced the protein levels of MMP3 and SDC4 (Fig. 5l-m).

Collectively, these findings demonstrate that mimic 0-4 effectively preserves cartilage ECM homeostasis in human OA models and highlight its potential as a chondroprotective therapeutic strategy, establishing a generalizable AI-driven paradigm for RNA therapeutic discovery.

## 3 Discussion

In this work, we presented RDiffusion, a conditional discrete diffusion model for comprehensive and programmable RNA sequence design. Unlike existing approaches that typically address isolated design tasks or rely on single-condition inputs, RDiffusion provides a unified framework capable of integrating diverse conditioning signals—including functional descriptions, RNA family types, secondary structures, tertiary structures, protein-binding and disease specifications—within a single generative architecture. This versatility enables RDiffusion to tackle a broad spectrum of RNA design challenges, ranging from unconditional sequence generation to highly constrained structure-guided and function-guided design.

Our experimental evaluation across multiple design tasks yielded several key insights. First, in functional RNA design for RNA family, 5’UTR MRL optimization and gRNA activity enhancement, RDiffusion consistently outperformed existing methods, achieving superior prediction accuracy and design success rates. The GRPO fine-tuning framework, combined with guidance model steering, proved particularly effective for optimizing sequences toward high-value functional properties while maintaining distributional fidelity. Second, in secondary structure-based design, RDiffusion demonstrated strong self-consistency and achieved high success rates on the challenging bpRNA TS0 benchmark, even without fine-tuning on the target dataset. Notably, RDiffusion generated sequences with substantially lower sequence similarity to wild-type sequences while maintaining structural fidelity, indicating genuine sequence novelty rather than training set memorization. The OpenKnot and Eterna score analyses further confirmed that RDiffusion-designed sequences exhibit folding properties comparable to—and in some cases exceeding—those of naturally occurring sequences. Third, in tertiary structure-based design on the RNASolo dataset, RDiffusion achieved significantly higher recovery rates than existing methods across short, medium, and long RNA sequences. The self-consistency TM-score and RMSD analyses revealed that RDiffusion-designed sequences refold into target structures with high accuracy, approaching the performance of wild-type sequences. The pLDDT scores from AlphaFold3 predictions further validated the structural rationality of RDiffusion-generated sequences. Fourth, in binding protein-based design, RDiffusion demonstrated strong performance in both single-protein and multi-protein targeting scenarios. The high success rates across 100 distinct binding proteins and 30 protein pairs underscore RDiffusion’s capability to navigate the complex trade-off between binding affinity and specificity, a central challenge in RNA aptamer and synthetic biology applications. Finally, unconditional generation experiments confirmed that RDiffusion produces structurally stable and diverse RNA sequences across a wide range of lengths, with short sequences achieving particularly high OpenKnot, Eterna, and diversity scores. The AlphaFold3-predicted pLDDT scores further supported the structural reasonability of unconditionally generated sequences.

To translate this computational framework into disease applications, we targeted OA as a prime paradigm, utilizing the RDiffusion to perform de novo design of novel miRNA sequences guided by a customized, data-driven seed selection and screening pipeline. Through the strategy, we identified the lead candidate mimic 0-4, which demonstrated superior chondroprotective efficacy against ECM dysregulation compared to its native miRNA sharing the same seed sequence. Mechanistically, the non-seed sequence optimization by RDiffusion enhanced target binding affinity toward SDC4, thereby downregulating its expression to effectively preserve ECM homeostasis and cartilage integrity. Crucially, this robust protective efficacy was consistently validated across clinical OA patient samples, reinforcing its translational relevance. Beyond providing a promising therapeutic lead for OA, this study establishes a generalizable AI-driven paradigm for RNA therapeutic discovery.

In conclusion, RDiffusion represents a significant advance in programmable RNA design, offering a unified, flexible, and powerful framework that integrates diverse conditioning signals to generate novel, structurally stable, and functionally optimized RNA sequences. We anticipate that RDiffusion will serve as a foundational tool for the RNA design community, accelerating both fundamental discovery and translational applications in RNA biotechnology.

## 4 Methods

### 4.1 Pretraining Data Preparation

The pretraining of RDiffusion requires a large-scale, high-quality dataset of RNA sequences annotated with both secondary structures and functional descriptions. To this end, we curated a comprehensive dataset from the RNAcentral database, which serves as a unified repository for non-coding RNA sequences from diverse organisms and functional classes.

#### 4.1.1 Data Extraction

We downloaded all RNA sequences from RNAcentral that are accompanied by two types of annotations: (i) predicted or experimentally determined secondary structures, and (ii) functional descriptions or Gene Ontology (GO) terms [102]. The secondary structure annotations provide crucial information about the spatial organization of nucleotides, which is essential for learning structure-function relation-ships. The functional descriptions offer textual summaries of biological roles, such as “ribozyme,” “riboswitch,” “tRNA,” “rRNA,” or “regulatory RNA,” enabling the model to incorporate functional context during training.

For each entry, we retained the following information:

- The raw RNA sequence represented as a string over the alphabet *{*A, U, C, G*}*.
- The secondary structure annotation in dot-bracket notation, where matched parentheses indicate base-paired regions and dots denote unpaired nucleotides.
- The functional description text, which may be a free-text sentence or a structured ontology term.
- Additional metadata including organism source, RNA family classification, and literature references where available.

#### 4.1.2 Redundancy Reduction via Sequence Similarity Clustering

To mitigate the effects of data redundancy and prevent overfitting to overrepresented sequence families, we performed a redundancy reduction step based on sequence similarity clustering. High sequence similarity between training examples can bias the model toward specific sequence patterns and reduce its generalization capability to novel sequences.

Specifically, we applied the MMseq2 tool [103] to cluster all extracted sequences at a similarity threshold of 90%. The clustering procedure is formally described as follows. Let *S* = {**s**_1_, **s**_2_, …, **s**_*N*_} denote the set of all extracted RNA sequences. For any two sequences **s**_*i*_ and **s**_*j*_, let sim(**s**_*i*_, **s**_*j*_) denote their sequence identity computed via global alignment. MMseq2 greedily clusters sequences such that:

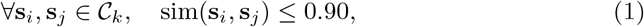

where *C*_*k*_ represents the *k*-th cluster, and the representative sequence for each cluster is the longest sequence within that cluster. All sequences with pairwise similarity exceeding 90% are grouped into the same cluster, and only the representative sequence (or a single randomly selected sequence from each cluster) is retained for the final dataset.

This clustering step reduced sequence redundancy by removing near-duplicate entries while preserving the diversity of the original data.

### 4.2 RDiffusion: A Discrete Diffusion Model for RNA Sequence Design

RDiffusion is a generative model based on discrete diffusion and transformer architecture, proposed for *de novo* RNA sequence design. The model learns to reconstruct masked tokens under varying corruption levels and can be conditioned on auxiliary information to steer generation toward sequences with desired functional properties.

#### 4.2.1 Training Phase

Let an RNA sequence be represented as a discrete token sequence **x** = (*x*_1_, *x*_2_, …, *x*_*L*_), where each token *x*_*i*_ belongs to a finite vocabulary *V* (e.g., {A, U, C, G}). During training, we first sample a mask ratio *λ ∼ u* (0, 1) uniformly from the interval [0, 1]. Subsequently, each position *i* in the sequence is independently masked with probability *λ*. The resulting corrupted sequence 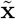 is obtained by replacing masked tokens with a special [MASK] token, while unmasked tokens retain their original values.

The model is additionally conditioned on auxiliary information **c**, which may encode structural constraints, functional annotations, or other task-specific requirements. The condition **c** is integrated into the model architecture via cross-attention or feature concatenation. The training objective is to maximize the log-likelihood of correctly predicting the original tokens at the masked positions:

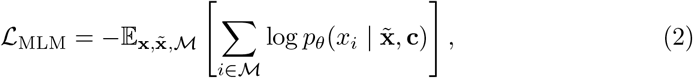

where *M* denotes the set of masked positions, and 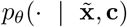 is the conditional distribution over the vocabulary *V* parameterized by the RDiffusion model with parameters *θ*. This objective, known as masked language modeling, trains the model to recover the original tokens given a partially observed input and conditioning signals. By exposing the model to a wide range of mask ratios *λ*, the model learns robust representations of RNA sequence logic that generalize across arbitrary levels of input corruption.

#### 4.2.2 Inference Phase

During inference, RDiffusion generates novel RNA sequences starting from an initial sequence that is either fully masked or partially masked. Generation proceeds in a stepwise denoising manner, where at each step a single masked position is selected and recovered. The process continues until all [MASK] tokens have been replaced, yielding a complete sequence.

Let **x**^(0)^ denote the initial sequence. In the fully masked setting, all positions are set to the [MASK] token; in the partially masked setting, a subset of positions may retain observed tokens as prior information. At each generation step *t* = 1, 2, …, *L*, the model performs the following operations:

1. **Reverse step**: The current partially masked sequence **x**^(*t−*1)^ is fed into the RDiffusion model together with the condition **c**. The model computes a probability distribution over the vocabulary *V* for each masked position:

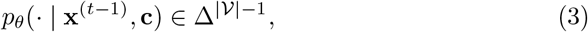

where Δ^|*V*|*−*1^ denotes the probability simplex over the vocabulary.
2. **Position selection**: A single masked position *i*_*t*_ is selected for recovery in the current step. The selection strategy can be random, deterministic (e.g., left-to-right), or based on model confidence. To ensure generation diversity, we adopt a random ordering sampled uniformly from all permutations of the masked positions.
3. **Token prediction**: For the selected position *i*_*t*_, the token is sampled from the predicted categorical distribution:

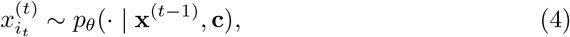

and the mask at position *i*_*t*_ is then removed.
4. **Sequence update**: The updated sequence is defined as:

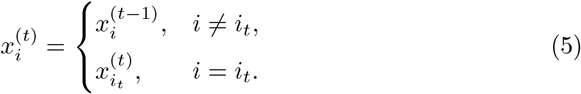

After exactly *L* steps, all masked positions have been recovered, yielding **x**^(*L*)^ as the final designed sequence. This iterative refinement procedure allows the model to leverage the context of already recovered tokens to inform subsequent predictions, thereby capturing long-range dependencies and complex sequence constraints.

To enable conditional generation of sequences with optimized functional properties, we further incorporate guidance from an auxiliary reward model. Let *R*(**x**) be a reward function (e.g., a separately trained predictor of mean ribosome load, MRL). The guided sampling distribution is defined as:

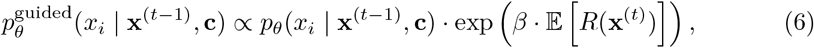

where **x**^(*t*)^ denotes the sequence after assigning token *x*_*i*_ to position *i*_*t*_, *β* is a temperature parameter controlling the strength of the guidance, and the expectation is taken over the stochasticity of future generation steps. In practice, this guidance can be approximated by treating the reward as a scoring function and reweighting the predicted probabilities accordingly. This guided inference scheme enables RDiffusion to generate sequences that not only respect the learned sequence distribution but also exhibit enhanced functional activity, such as high translational efficiency or improved stability.

### 4.3 Condition Injector Module

To enable flexible and comprehensive control over the generation process, RDiffusion incorporates a Condition Injector module that integrates multiple types of conditioning information. Each condition type is encoded into a latent representation and subsequently fused with the RNA sequence embeddings through addition. Formally, let **E** ∈ ℝ^*B×L×D*^ denote the initial RNA sequence embeddings, where *B* is the batch size, *L* is the sequence length, and *D* is the hidden dimension. The Condition Injector processes four distinct categories of conditions, as detailed below.

#### 4.3.1 Textual RNA Description Condition

The first condition type consists of natural language descriptions of the desired RNA functional properties. Given a text description of length *L*_*t*_, represented as 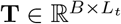, we employ a pretrained BioBERT [104] model to encode the text into contextualized embeddings. The BioBERT encoder, denoted as *f*_BioBERT_, maps the input text to a sequence of token embeddings:

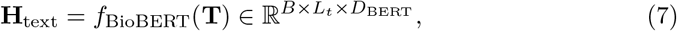

where *D*_BERT_ is the output dimension of BioBERT. To obtain a fixed-length global representation, we apply mean pooling over the sequence length dimension:

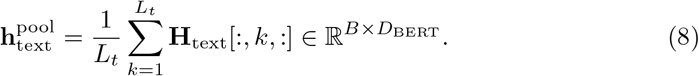

This pooled representation is then projected into the RNA latent space using a learnable projection function 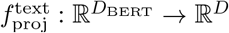:

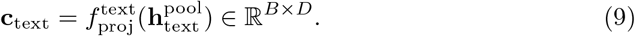

Finally, the textual condition is broadcast across the sequence length dimension and added to the RNA embeddings:

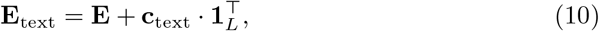

where **1**_*L*_ *∈* ℝ^*L*^ is an all-ones vector.

#### 4.3.2 Functional Category Condition

The second condition type encompasses discrete or continuous functional annotations, such as desired activity levels, stability metrics, or regulatory categories. Let **f** *∈* ℝ^*B×*1^ denote the raw functional condition (e.g., a scalar activity score or a categorical label). We employ an embedding module 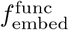, which can be either a standard embedding lookup table for discrete categories or a linear projection for continuous values:

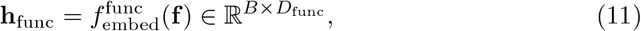

where *D*_func_ is the intermediate embedding dimension. A subsequent projection function 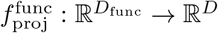 maps this representation into the RNA latent space:

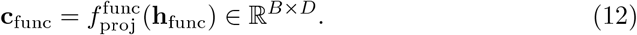

Similar to the textual condition, **c**_func_ is broadcast across the sequence length and added to the RNA embeddings:

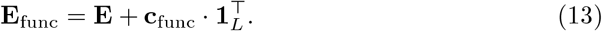

#### 4.3.3 Secondary Structure Condition

The third condition type encodes the RNA secondary structure at the sequence level. Given a secondary structure annotation (e.g., in dot-bracket notation or numerical format), we convert it into a per-nucleotide feature vector of dimension 10, following the procedure outlined in Algorithm 1. The resulting structure feature tensor is denoted as **S** *∈* ℝ^*B×L×*10^.

These 10-dimensional per-nucleotide features capture local structural context, including pairing probability, nesting depth, and loop type information. The structure features are then projected into the RNA latent space using a sequence projection function 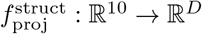:

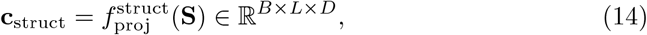

##### Algorithm 1 Structure to Feature Vector Conversion

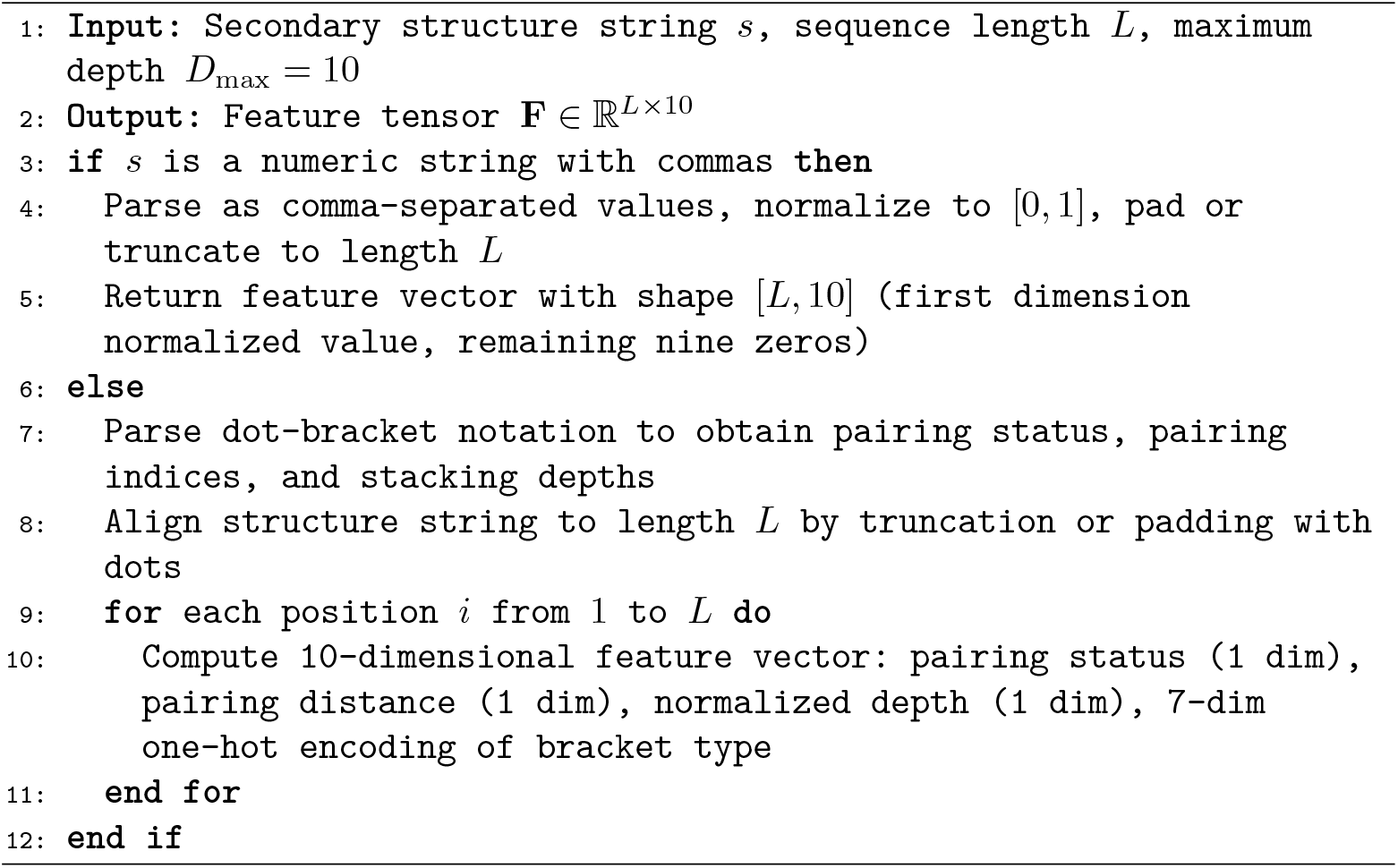

where 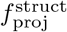 is implemented as a multi-layer perceptron applied independently to each position. Unlike the previous two conditions, the secondary structure condition provides position-specific information and is therefore added directly to the RNA embeddings in a position-wise manner:

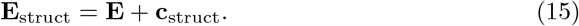

#### 4.3.4 Tertiary Structure Condition

The fourth condition type encodes the RNA tertiary (3D) structure. We first construct a geometric graph representation from the 3D coordinates of the RNA backbone. Following the featurization strategy used in gRNAde, we adopt a 3-bead coarse-grained representation comprising the P, C4’, and N1 (pyrimidine) or N9 (purine) atoms for each nucleotide. For each nucleotide *i*, we compute its centroid coordinate **x**_*i*_ *∈* ℝ^3^ as the centroid of the three bead atoms. The resulting point cloud is used to construct a *k*-nearest neighbor graph, where each node is connected to its 32 nearest neighbors based on Euclidean distance ∥**x**_*i*_*−* **x**_*j*_ ∥_2_.

Node features are initialized with geometric descriptors, including forward and reverse backbone unit vectors, as well as distances, angles, and torsions between the C4’ atom and its corresponding P and N1/N9 atoms. Edge features for each directed edge from node *j* to *i* include the unit vector ^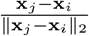^, the distance ∥**x**_*j*_ *−* **x**_*i*_∥ _2_ encoded via 32 radial basis functions, and the backbone distance *j i* encoded via 32 sinusoidal positional encodings.

This geometric graph is then processed by a Graph Neural Network (GNN) to obtain per-nucleotide structural embeddings. Let *G* = (*V, E*) denote the constructed graph with node features 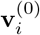 and edge features **e**_*ji*_. The GNN, denoted as *f*_GNN_, performs iterative message passing:

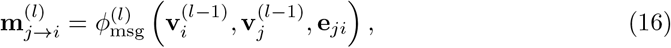

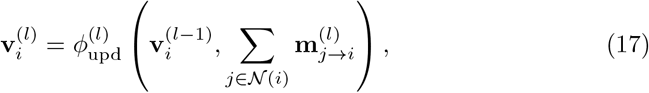

where 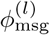 and 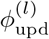 are learnable message and update functions at layer *l*, and *N* (*i*) denotes the set of neighbors of node *i*. After *L*_GNN_ layers, the final node embeddings are obtained as 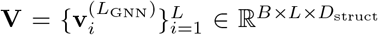, where *D*_struct_ is the structural feature dimension.

Finally, a sequence projection function 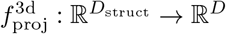 maps these structural embeddings into the RNA latent space:

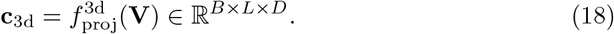

The tertiary structure condition is then fused with the RNA embeddings via position-wise addition:

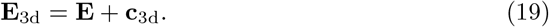

#### 4.3.5 Condition Fusion

In practice, multiple condition types may be available simultaneously. The over-all conditioned RNA embeddings are obtained by sequentially applying all available condition additions:

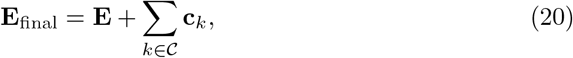

where *C* denotes the set of active condition types (textual, functional, secondary structure, and/or tertiary structure), and each **c**_*k*_ is appropriately broadcast or position-wise as described above. This additive fusion mechanism ensures that the model can flexibly incorporate diverse sources of information while maintaining a simple and interpretable conditioning architecture.

### 4.4 Guidance Model for Conditional Generation

To enable controllable generation of RNA sequences with desired functional properties, we introduce a guidance mechanism that leverages an auxiliary predictive model. This approach allows RDiffusion to steer the generation process toward sequences that exhibit specific characteristics, such as high MRL or enhanced structural stability, without requiring retraining of the diffusion model itself.

#### 4.4.1 Training of the Guidance Model

Let **x** = (*x*_1_, *x*_2_, …, *x*_*L*_) denote an RNA sequence, and let *y* ∈ *Y* be the corresponding label or property value of interest (e.g., a scalar activity score or a categorical functional class). We train a guidance model *g*_*ϕ*_, parameterized by *ϕ*, to predict the label *y* from the RNA sequence **x**. The guidance model can be implemented as any sequence-to-property architecture, such as a convolutional neural network, a transformer encoder, or a pretrained language model fine-tuned for regression or classification.

Given a training dataset 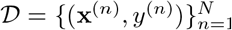 of RNA sequences paired with their measured or predicted properties, the guidance model is trained to minimize a loss function *L*_guide_ that quantifies the discrepancy between the predicted and ground-truth labels:

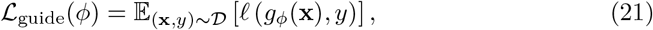

where *l*(*·,·*) is a task-appropriate loss function. For regression tasks (e.g., predicting MRL values), we typically employ the mean squared error loss:

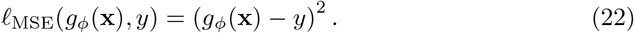

For classification tasks (e.g., predicting functional categories), we employ the cross-entropy loss:

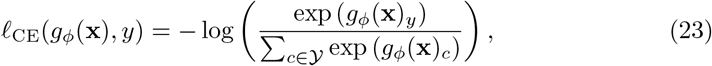

Where *g*_*ϕ*_(**x**)_*c*_ denotes the logit corresponding to class *c*. The parameters *ϕ* are updated via standard gradient-based optimization:

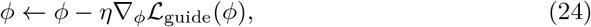

where *η* is the learning rate. Upon convergence, the trained guidance model provides a differentiable mapping from RNA sequences to property predictions, which can be used to guide the diffusion generation process.

#### 4.4.2 Guidance During RDiffusion Inference

During the inference phase of RDiffusion, we aim to generate RNA sequences that not only follow the learned sequence distribution but also exhibit high predicted property values according to the guidance model. Let **x**^(*t−*1)^ denote the partially masked sequence at generation step *t −* 1, and let **p**_*θ*_(*·*| **x**^(*t−*1)^, **c**) be the probability distribution over the vocabulary *V* predicted by RDiffusion for each masked position, parameterized by *θ*. For a selected masked position *i*_*t*_, the raw logits before softmax are denoted as **z** *∈* ℝ^|*V*|^, where:

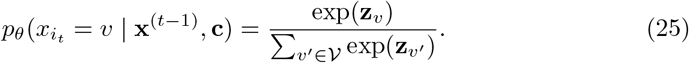

To bias the generation toward sequences with favorable properties while maintaining end-to-end differentiability, we treat the guidance model’s input as a continuous relaxation of the discrete token distribution. Specifically, instead of sampling a discrete token *v* and constructing a hard one-hot sequence 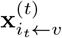, we construct a continuously relaxed sequence 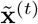 where the token at position *i*_*t*_ is represented by the softmax distribution itself, and all other positions remain as their current (possibly also soft) representations. Formally, let 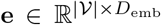 be the token embedding matrix. The continuous embedding for position *i*_*t*_ is given by:

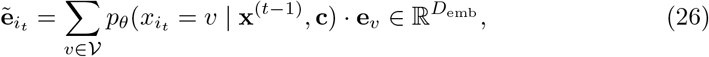

which is a convex combination of token embeddings weighted by the predicted probabilities. The continuously relaxed sequence 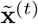 is then fed into the guidance model *g*_*ϕ*_, which predicts a label 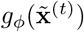 in a fully differentiable manner.

We define a guidance loss 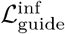 that encourages the model to assign high probability to tokens that lead to desirable property values:

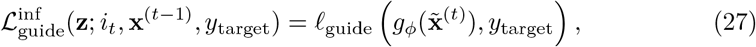

where *y*_target_ is a target property value (e.g., a high MRL score for maximization tasks), and *l*_guide_ is a loss function that penalizes deviations from the target. For maximization tasks, we typically set *y*_target_ to a high constant value and use a negative reward formulation, e.g., *l*_guide_(*ŷ, y*_target_) =*− ŷ* to encourage higher predicted values.

Importantly, because 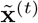 depends smoothly on the logits **z** through the softmax function, the gradient 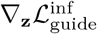 can be computed directly via standard backpropagation without requiring REINFORCE or other gradient estimators. This enables efficient and stable guidance of the diffusion process.

In practice, to obtain a valid discrete sequence at the end of generation, we can optionally use the straight-through Gumbel-Softmax estimator [105] during training of the guidance loop, where the forward pass uses a discrete sample while the backward pass uses the continuous softmax relaxation. However, for inference-time guidance as described here, the continuous relaxation alone is sufficient to produce meaningful gradients.

In practice, we approximate the expectation by considering the current predicted distribution. The gradient of the guidance loss with respect to the logits **z** is computed via backpropagation through the guidance model. Specifically, we compute:

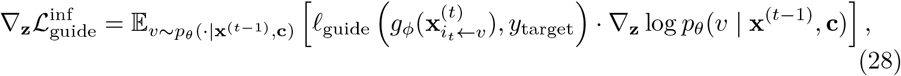

which can be efficiently estimated using the REINFORCE algorithm [106] or by reparameterization when the guidance model is differentiable. The guided logits are then obtained by performing one or more gradient ascent steps on the original logits:

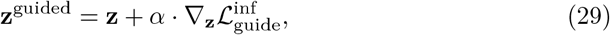

where *α* is a step size hyperparameter controlling the strength of the guidance. Alternatively, a simpler and computationally efficient approach is to directly modify the sampling distribution by weighting the logits with the guidance model’s output:

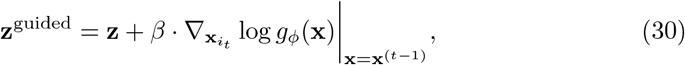

where the gradient is taken with respect to the input token at position *i*_*t*_, approximated via finite differences or through a differentiable embedding layer.

After obtaining the guided logits **z**^guided^, the updated probability distribution is computed as:

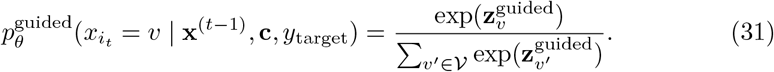

The token for position *i*_*t*_ is then sampled from this guided distribution:

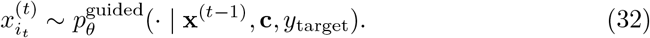

This guided sampling procedure is repeated at each generation step, allowing the model to iteratively refine the sequence while being continuously steered toward the desired property space.

#### 4.4.3 Algorithm Summary

The overall guided generation process is summarized in Algorithm 2. The key advantage of this approach is that the guidance model is trained separately and can be easily swapped or combined, enabling flexible control over multiple properties without modifying the underlying diffusion model.

##### Algorithm 2 Guided Generation with RDiffusion

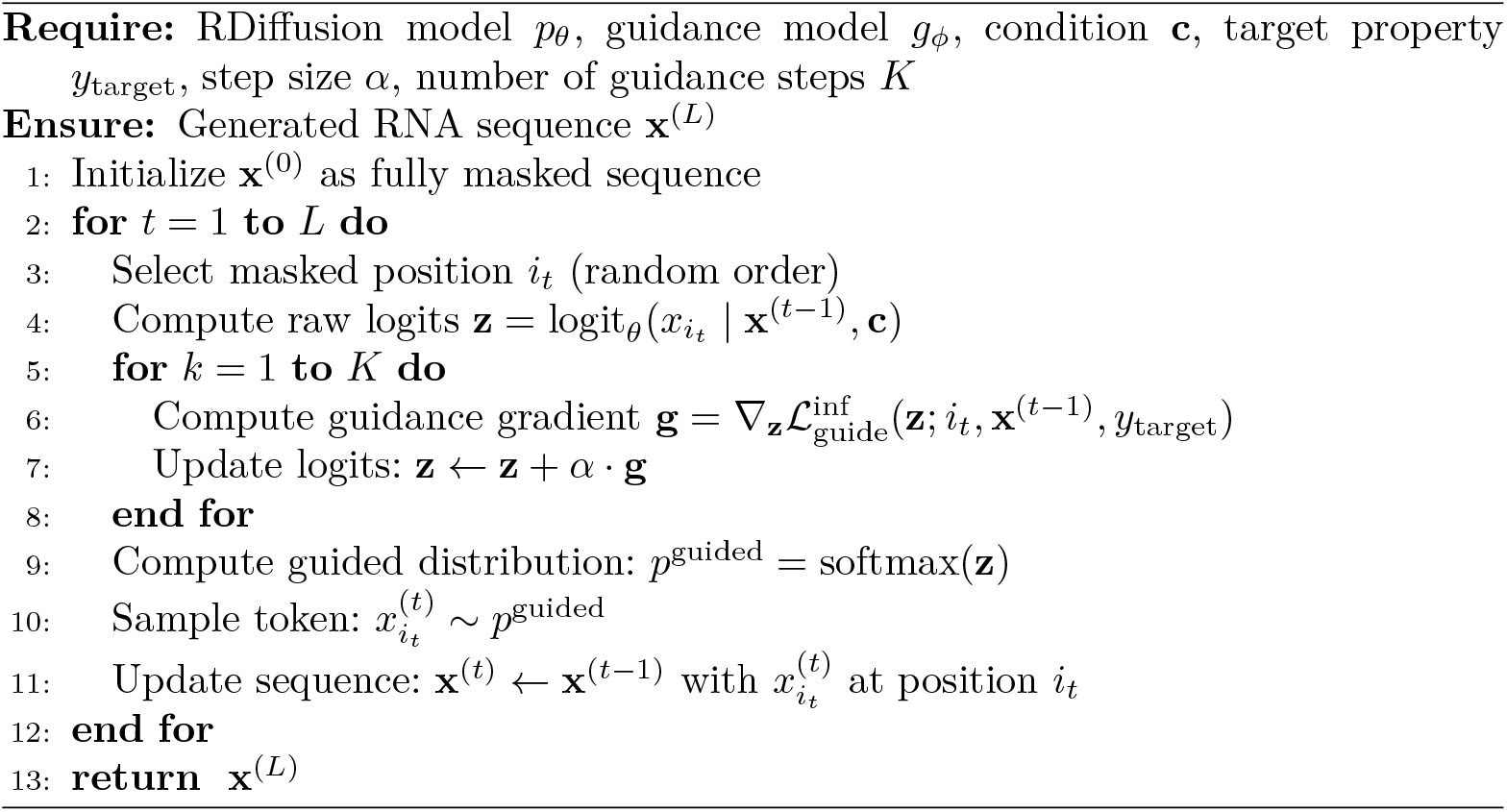

This guidance mechanism provides a flexible and efficient way to control the properties of generated RNA sequences, enabling targeted design of molecules with optimized functional characteristics.

### 4.5 Group Reward Policy Optimization for RDiffusion Fine-Tuning

After obtaining a pretrained guidance model *g*_*ϕ*_ that serves as a differentiable reward model, we further fine-tune the RDiffusion model using Group Reward Policy Optimization (GRPO). This training paradigm treats the pretrained RDiffusion as both the policy model and the reference model, enabling it to optimize toward generating sequences with high reward while maintaining distributional constraints.

#### 4.5.1 Preliminaries and Notations

Let *π*_*θ*_ denote the policy model (the RDiffusion model being fine-tuned) parameterized by *θ*, and let *π*_ref_ denote the reference model (the pretrained RDiffusion with frozen parameters). Given a conditioning signal **c** (which may include functional requirements, structural annotations, or target property specifications) and a partially masked sequence, the policy model generates a probability distribution over masked tokens. For a training batch, we sample a set of *G* sequences {**x**^(1)^, **x**^(2)^, …, **x**^(*G*)^} from the policy model, where each sequence is sampled from the probability distribution.

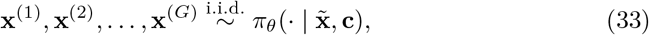

where 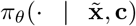 denotes the conditional distribution over complete RNA sequences defined by the policy model.

The reward for each generated sequence is computed using the guidance model *g*_*ϕ*_:

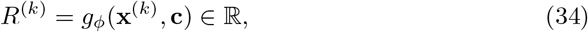

where higher reward values indicate sequences that better satisfy the desired functional properties (e.g., higher predicted MRL or improved structural stability).

#### 4.5.2 Group Reward Policy Optimization Objective

The GRPO algorithm optimizes the policy model by comparing each generated sequence against a group baseline computed from the *G* samples. The advantage for the *k*-th sequence is defined as:

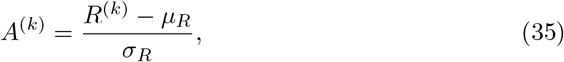

where 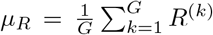 and 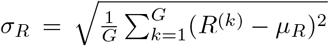 are the mean and standard deviation of the rewards within the group, respectively. This group-based normalization reduces variance in policy gradient estimation without requiring a separate value network.

The GRPO loss encourages the policy to assign higher probability to sequences with positive advantages while discouraging those with negative advantages. It is defined as:

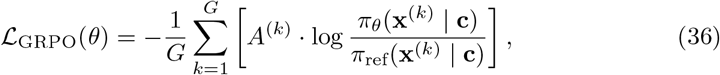

where the ratio 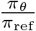 ensures that the policy does not deviate too far from the reference model, providing a form of trust region constraint.

#### 4.5.3 Training Procedure with Masked Language Modeling

During GRPO training, the RDiffusion model continues to follow the discrete diffusion training paradigm. Specifically, for each RNA sequence **x** in the training batch, we first sample a random mask ratio *λ ∼ U* (0, 1) and independently mask each position with probability *λ*, yielding a corrupted sequence 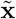. The model then predicts the original tokens at masked positions:

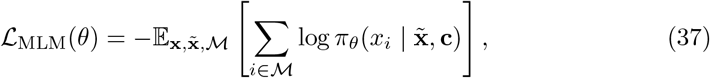

where *M* denotes the set of masked positions. This objective ensures that the model retains its ability to recover masked tokens accurately.

#### 4.5.4 Cross-Entropy Loss for Sequence Consistency

To maintain consistency with the original sequence distribution and prevent catastrophic forgetting, we introduce a cross-entropy loss between the policy model’s predictions and the original training data distribution. Let **x**^orig^ denote the original uncorrupted sequence. The cross-entropy loss is defined as:

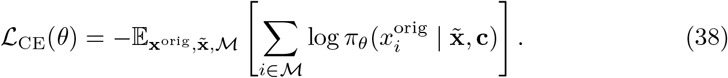

This loss encourages the model to preserve the original sequence identity at masked positions, thereby maintaining fidelity to the training distribution.

#### 4.5.5 Reference Loss for Distributional Regularization

In addition to the cross-entropy loss, we employ a reference loss that directly regularizes the Kullback-Leibler (KL) divergence [107] between the policy and reference model predictions. For each masked position *i ∈ M*, let 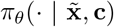 and 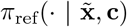 denote the probability distributions over the vocabulary predicted by the policy and reference models, respectively. The reference loss is defined as:

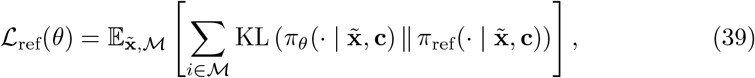

where the KL divergence is computed as:

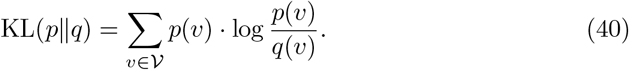

This loss penalizes large deviations from the reference model, ensuring that the fine-tuned policy remains close to the pretrained distribution while still adapting to the reward signal.

##### Algorithm 3 GRPO Training for RDiffusion

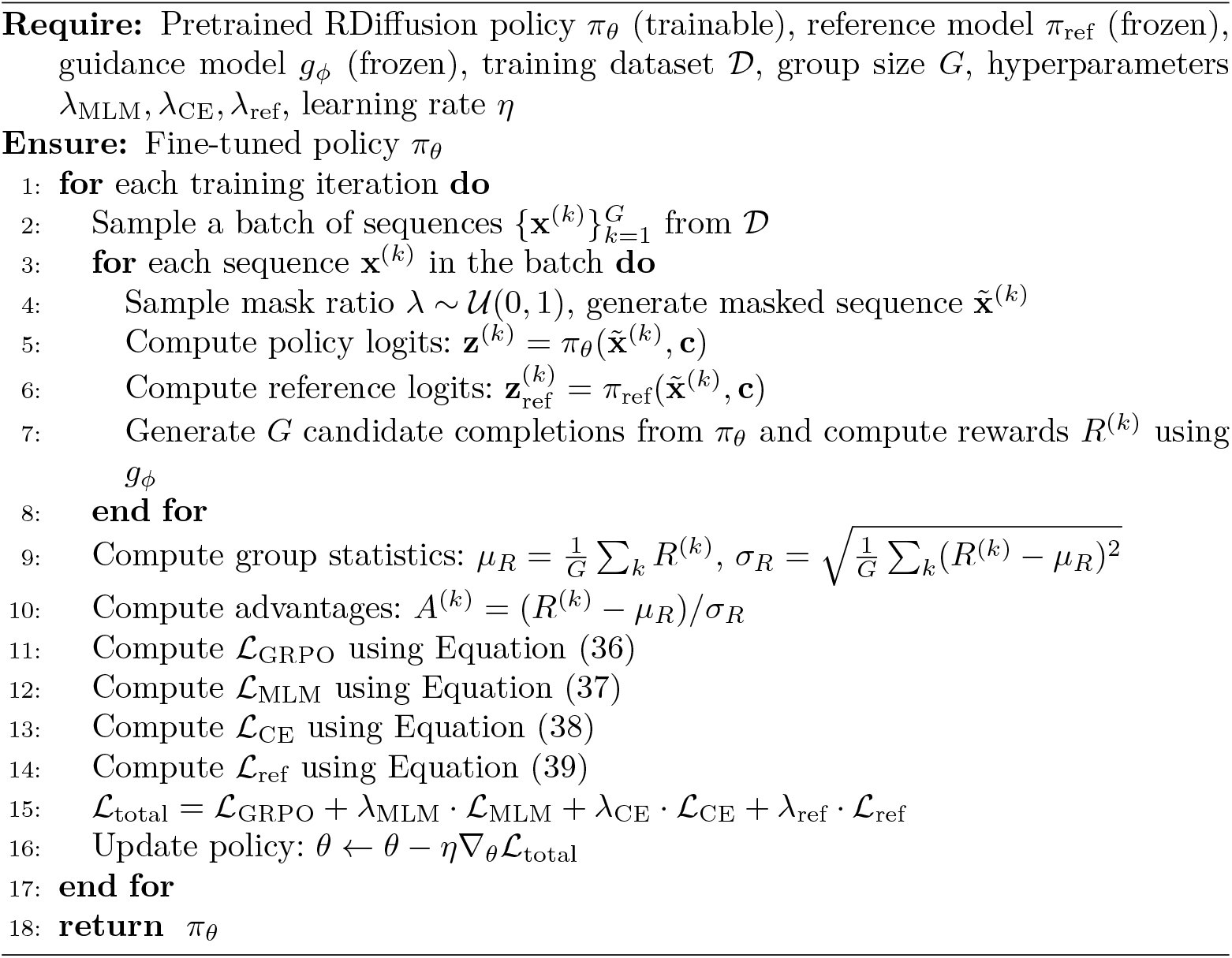

#### 4.5.6 Total Training Objective

The overall training objective for RDiffusion under GRPO is a weighted combination of the GRPO loss, the masked language modeling loss, the cross-entropy loss, and the reference loss:

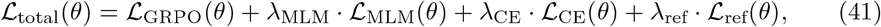

where *λ*_MLM_, *λ*_CE_, and *λ*_ref_ are hyperparameters controlling the relative importance of each loss term. In practice, we set *λ*_MLM_ = 1.0, *λ*_CE_ = 0.1, and *λ*_ref_ = 0.01 to balance reward optimization with distributional constraints.

The parameters *θ* are updated via gradient descent:

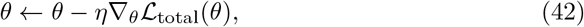

where *η* is the learning rate.

#### 4.5.7 Algorithm Summary

The complete GRPO training procedure for RDiffusion is summarized in Algorithm 3.

This GRPO-based fine-tuning framework enables RDiffusion to iteratively improve its sequence design capability toward high-reward functional objectives while maintaining distributional fidelity to the original training data.

### 4.6 Training Details

This section provides comprehensive details on the training configuration for both the pretraining stage and subsequent downstream fine-tuning stages. All experiments were implemented using the PyTorch framework and conducted on NVIDIA A100 GPUs.

#### 4.6.1 Pretraining Configuration

For the pretraining phase, we optimized the RDiffusion model using the AdamW optimizer [108] with a learning rate of 1*×* 10^*−*4^. The training was performed with a global batch size of 16, combined with gradient accumulation over 8 batches to effectively simulate a larger batch size while respecting memory constraints. The learning rate was scheduled using a cosine decay schedule [109].

The pretraining was distributed across 8 NVIDIA A100 GPUs using data parallelism. The total training duration was approximately 240 hours. The complete pretraining hyperparameters are summarized in Table 1.

**Table 1.** Pretraining hyperparameters for RDiffusion.

| **Hyperparameter** | **Value** |
|---|---|
| Optimizer | AdamW |
| Learning rate | 1 × 10<sup>−4</sup> |
| Learning rate schedule | Cosine decay |
| Batch size (per device) | 16 |
| Gradient accumulation steps | 8 |
| Effective batch size | 128 |
| Number of GPUs | 8 |
| GPU type | NVIDIA A100 |
| Training duration | 240 hours |

#### 4.6.2 Downstream Fine-Tuning Configuration

For downstream task fine-tuning, we adopted a modified training configuration while keeping most hyperparameters consistent with the pretraining setup. The fine-tuning configuration differs in the following aspects: Gradient accumulation: 1, Learning rate schedule: one-cycle schedule, Training device: 1 NVIDIA A100 GPU All other hyper-parameters remain identical to the pretraining configuration, including the optimizer (AdamW), base learning rate (1 *×* 10^*−*4^), and per-device batch size (16).

#### 4.6.3 Implementation Details

All models were implemented using PyTorch with the Pytorch lightning library [110] for efficient training and deployment. Mixed precision training using automatic mixed precision (AMP) was enabled for both pretraining and fine-tuning to accelerate computation and reduce memory usage. Gradient clipping with a maximum norm of 1.0 was applied to prevent gradient explosion.

### 4.7 Clinical Samples and Ethics Statement

Human OA cartilage samples were obtained from patients undergoing total knee arthroplasty at Zhongshan Torch Development Zone People’s Hospital. Written informed consent was obtained from all participants and all samples were anonymized before analysis. The study was approved by the Ethics Committee of Zhongshan Torch Development Zone People’s Hospital (Approval No. 2025-209).

### 4.8 Cell Culture and Treatment

Chondrosarcoma cell line SW1353 maintained in DMEM with 10% fetal bovine serum (26140079, Gibco Life Technologies, Oshima, New York) and penicillin-streptomycin (15140122, Gibco Life Technologies, Oshima, New York), were treated with IL-1*β* (10 ng/mL, 10139-HNAE, Sino Biological Inc, Beijing, China) for 48 h.

Primary human chondrocytes were isolated as previously described [111]. Briefly, articular cartilage tissues were minced into approximately 1 mm^3^ fragments and washed three times with PBS containing 100 U/mL penicillin and 100 *µ*g/mL streptomycin. The fragments were initially digested with 0.25% trypsin (27250-018, Invitrogen, Carlsbad, CA, USA) in a 37^*°*^C water bath for 30 min, and the digestion was terminated by the addition of FBS. After three subsequent washes with PBS, the tissue was further digested with 0.2% type II collagenase (40508ES60, Yeasen Biotechnology, Shanghai, China) overnight at 37^*°*^C. The resulting cell suspension was passed through a 70 *µ*m cell strainer and centrifuged at 1000 rpm for 10 min. The isolated chondrocytes were then resuspended and seeded into culture dishes for further experiments.

Mimic, siRNA SDC4 and ADAM10 was purchased from BiOligo Biotechnology, Shanghai, China. si-SDC4 (sense (5^*′*^–3^*′*^): GCUAUGACCUGGGCAAGAA, anti-sense (5^*′*^–3^*′*^): UUCUUGCCCAGGUCAUAGC); si-ADAM10 (sense (5^*′*^–3^*′*^): GCTGTGCA-GATCATTCAGTAT, anti-sense (5^*′*^–3^*′*^): ACUGAAUGAUCUGCACAGC).

### 4.9 Explant Culture

Human OA cartilage explants (about 4 *×* 4 mm) were placed in 24-well cell plates, after the explants were cultured in normal high glucose medium for 48 hours to adapt to the stable state of in vitro culture environment, they were stimulated with 10 ng/mL IL-1*β* for 48 hours and transfection mimic.

### 4.10 Western Blot

Protein samples were processed with SDS loading buffer and boiled to denature the proteins. Then, the denatured samples were separated based on size using 10%–15% SDS-polyacrylamide gel electrophoresis, followed by transferring the proteins of interest to a membrane. The membrane was blocked with 5% milk or BSA solution for 1 h to prevent nonspecific antibody binding. Then, the membrane was incubated with the appropriate primary antibodies at 4^*°*^C overnight, followed by incubation with a secondary antibody at room temperature for 1 h. Finally, the membrane was stained with ECL detection buffer (1705061, Bio-Rad, Hercules, CA, USA) and imaged by a ChemiDoc system. Primary antibodies included *β*-actin (4970, Cell Signaling Technology, Trask Lane Danvers, MA, USA), a senescence marker antibody sampler kit (56062T, Cell Signaling Technology, Trask Lane Danvers, MA, USA) containing p21, MMP3, SOX9 (A19710, ABclonal, Wuhan, China), MMP13 (50497, genuinbiotech, Hefei, China), SDC4 (11820-1-AP, Proteintech, Wuhan, China).

### 4.11 RNA Extraction and Quantitative Real-time Polymerase Chain Reaction

Total RNA was extracted using 200 *µ*L of TRIzol reagent (9019, TaKaRa, Japan). One microgram of the total RNA was transcribed into cDNA in 20 *µ*L reactions using the PrimeScript RT Master Mix (RR036, TaKaRa, Japan) and then diluted to 300 *µ*L, which was used in subsequent real-time qPCR reactions. Real-time qPCR was performed using the SYBR Green qPCR Kit (B21702, Bimake, Houston, TX, USA) on an Agilent Mx3000P system (Agilent Technologies, Santa Clara, CA, USA) as specified by the manufacturer. All reactions were performed in triplicate. Primer sequences were synthesized by Saiheng Biological Shanghai, China. The primer sequences: CCL2 (sense (5^*′*^ to 3^*′*^): AGAATCACCAGCAGCAAGT-GTCC, anti-sense (5^*′*^–3^*′*^): TCCTGAACCCACTTCTGCTTGG); MMP13 (sense (5^*′*^ to 3^*′*^): CCTTGATGCCATTACCAGTCTCC, anti-sense (5^*′*^–3^*′*^): AAACAGCTC-CGCATCAACCTGC); MMP3 (sense (5^*′*^ to 3^*′*^): CACTCACAGACCTGACTCG-GTT, anti-sense (5^*′*^–3^*′*^): AAGCAGGATCACAGTTGGCTGG); SOX9 (sense (5^*′*^ to 3^*′*^): AGGAAGCTCGCGGACCAGTAC, anti-sense (5^*′*^–3^*′*^): GGTGGTCCTTCTTGT-GCTGCAC); SDC4 (sense (5^*′*^ to 3^*′*^): GAGTGAGGATGTGTCCAACAAGG, anti-sense (5^*′*^–3^*′*^): GGTACATGAGCAGTAGGATCAGG). The data were averaged and normalized to the expression level of glyceraldehyde-3-phosphate dehydrogenase (*β*-actin).

### 4.12 Senescence-associated *β*-galactosidase (SA-*β*-gal) Staining

SA-*β*-gal staining was performed using a Senescence *β*-Galactosidase Staining Kit (C0602, Beyotime, Haimen, China) according to the manufacturer’s instructions. Cell samples were fixed with the fixation solution from the kit for 15 min. After three rinses with PBS for 5 min, the cells were incubated with freshly prepared SA-*β*-Gal staining solution overnight in a 37^*°*^C humidified chamber. The next day, the cells were washed three times in PBS for 5 min at room temperature and observed with a microscope.

### 4.13 Safranin O and Immunohistochemistry (IHC) Staining

The tissues were fixed in 4% paraformaldehyde and embedded in paraffin. Safranin O staining was performed. For immunohistochemical detection of MMP3 and SDC4 expression, sections were deparaffinized and rehydrated, endogenous peroxidase was quenched with 3% aqueous hydrogen peroxide for 15 min. Then, the sections were blocked with 5% BSA (B2064-10G, Sigma-Aldrich, St. Louis, MO, USA) in Tris-buffered saline for 1 h and incubated with the respective primary antibodies overnight at 4^*°*^C. After PBS washes, the secondary antibody (abs20002A, Absin Bioscience, Shanghai, China) was used. After incubation for 30 min, 3,3^*′*^-diaminobenzidine tetrahydrochloride was added for 10 min at room temperature. Finally, the slides were stained with hematoxylin and examined under a microscope.

### 4.14 Dual-luciferase Reporter Assay

HEK293T cells were seeded in 96-well plates and cultured in high-glucose Dulbecco’s modified Eagle’s medium supplemented with 10% fetal bovine serum and 1% penicillin–streptomycin at 37 ^*°*^C in a humidified incubator with 5% CO_2_. When cells reached approximately 70–80% confluence, co-transfection was performed using Lipo-fectamine 3000 (Invitrogen, Thermo Fisher Scientific) according to the manufacturer’s instructions.

Briefly, cells were co-transfected with a luciferase reporter plasmid containing the SDC4 3^*′*^untranslated region (3^*′*^UTR) (constructed by MiaoLingPlasmid, Wuhan, China) together with mimic 0-4 or the corresponding native miRNA (hsa-miR-140-3p). Six hours after transfection, the medium was replaced with fresh complete DMEM containing 10% FBS, and cells were cultured for an additional 48 h.

Luciferase activity was measured using the KeyTec®Firefly–Renilla Luciferase Detection Kit (A2000900N, VKEY Biotechnologies, Shanghai, China) according to the manufacturer’s protocol. Briefly, cells were washed twice with ice-cold phosphate-buffered saline (PBS), followed by the sequential addition of Firefly and Renilla luciferase detection reagents (50 *µ*L each). After incubation with gentle shaking (400 rpm, 10 min) following each reagent addition, Firefly and Renilla luminescence were measured using a microplate luminometer.

Relative luciferase activity was calculated by normalizing Firefly luciferase activity to Renilla luciferase activity (Firefly/Renilla). All experiments were performed independently at least three times.

### 4.15 RNA-seq Analysis for Different Groups of Chondrocytes

Sequencing analysis was performed on two groups: the IL-1*β* treatment group, and the 0-4+IL-1*β* co-treatment group. The sequencing service was completed by Shanghai Majorbio Bio-pharm Technology.

RNA integrity and concentration were assessed using an Agilent 2100 Bioanalyzer, and samples with an RNA integrity number (RIN) greater than 7.0 were used for library preparation. Poly(A)+ RNA was enriched using oligo (dT) magnetic beads, fragmented, and reverse-transcribed into cDNA. Sequencing libraries were prepared using a standard Illumina library preparation kit following the manufacturer’s protocol and sequenced on an Illumina NovaSeq 6000 platform to generate paired-end 150-bp reads.

Raw sequencing reads were subjected to quality control by removing adaptor sequences, low-quality reads, and reads containing excessive ambiguous nucleotides. Clean reads were aligned to the human reference genome using HISAT2, and gene expression levels were quantified with StringTie. Gene expression abundance was normalized as transcripts per million for visualization, whereas raw read counts were used for differential expression analysis.

Differentially expressed genes between the IL-1*β* group and the 0-4 + IL-1*β* group were identified using the DESeq2 package in R. Genes with an adjusted *P* value *<* 0.05 and an absolute log_2_ fold change *>* 1 were considered significantly differentially expressed. Hierarchical clustering analysis of DEGs was performed based on normalized expression values using Euclidean distance and complete linkage clustering. Heatmaps were generated using the R package pheatmap to visualize gene expression patterns across samples.

Kyoto Encyclopedia of Genes and Genomes pathway enrichment analysis was performed using the clusterProfiler package in R. Pathways with an adjusted *P* value *<* 0.05 were considered significantly enriched.

### 4.16 Statistical Analysis

Statistical analyses per experiment are indicated in figure legends. Statistical tests were performed in GraphPad Prism 9.0 using Student’s two-tailed unpaired *t* test for pairwise comparisons, one-way or two-way analysis of variance (ANOVA) for multiple comparisons. A *p* value less than 0.05 was considered significant.

## 5 Data Availability

The pretraining dataset of RFuncData is collected from the RNACentral database. The Rfam dataset is accessible at https://rfam.org/. The bpRNA-1m dataset is obtained from https://bprna.cgrb.oregonstate.edu/. The RNASolo dataset can be downloaded from https://rnasolo.cs.put.poznan.pl/archive. The protein-RNA binding datasets can be accessed at https://zhanglabnet.oss-cn-beijing.aliyuncs.com/prismnet/data/clip_data.tgz. The datasets for 5’UTR and gRNA design tasks can be downloaded from https://doi.org/10.1038/s41587-019-0164-5, https://github.com/pjsample/human_5utr_modeling, and https://www.ncbi.nlm.nih.gov/geo/query/acc.cgi?acc=GSM3130443, as well as from https://doi.org/10.1038/s41587-022-01213-5 and https://github.com/broadinstitute/adapt-seq-design/tree/main/data/CCF-curated, respectively. We also provide the processed test datasets and source data used in this work at https://modelscope.cn/datasets/wj1006/RDiffusion_test_data and https://doi.org/10.6084/m9.figshare.32249322 [112], respectively.

## 6 Code Availability

We provide the source code in https://github.com/A4Bio/RDiffusion/ under MIT license. This repository also contains detailed instructions and tutorials for RDiffusion’s applications to reproduce our results. The specific version of the code associated with this publication is archived in Zenodo and is accessible via https://doi.org/10.5281/zenodo.20159683 [113].

## 7 Acknowledgements

We thank Dr. Jing Wang from Zhongshan Torch Development Zone People’s Hospital for kindly providing the clinical OA patient samples. This work was supported by the National Natural Science Foundation of China Project (No.624B2115) and the Zhejiang Key Laboratory Project (2024E10001). This work is funded by the Innovative Drug Research and Development National Science and Technology Major Project (Grant No. 2025ZD1804100) and the Strategic Priority Research Program of the Chinese Academy of Sciences (Grant No. XDB1060000).

## 8 Author Contributions

Jue Wang conceived the idea. Jintong Dong collected and processed the pretraining, RNASolo, and Rfam datasets. Jue Wang and Jintong Dong pretrained the RDiffusion. Jue Wang finished the Rfam, secondary structure, protein-RNA-binding and unconditional tasks; Jintong Dong finished the tertiary structure task; Tianhao Li collected the datasets and finished the 5’UTR and Cas13 guide RNA design tasks. Ying Dong and Jia Li designed the overall framework for screening and validating AI-predicted miRNAs in disease models. Lan Yang performed the biological experiments and conducted the data analysis. Jue Wang drafted the manuscript; Lan Yang, Ying Dong, and Jia Li. contributed to the writing of the experimental validation section. Jin-tong Dong, Tianhao Li, Lan Yang, Jianwei Yin, Jintao Chen, Ying Dong, Jia Li, and Cheng Tan helped in revising the manuscript. Jue Wang, Jintong Dong, and Tianhao Li prepared the code and released it on GitHub. Jintao Chen, Ying Dong, Jia Li, and Cheng Tan supervised the project.

## Footnotes

1 The Hamming distance is equal to the number of characters that differ between two strings of equal length at the same positions.

